# Partial resistance to HDAC inhibitors in FAPs of dystrophic muscles at late stages of disease is associated to epigenetic and transcriptional features of cellular senescence

**DOI:** 10.1101/2021.04.26.441412

**Authors:** S. Consalvi, L. Tucciarone, E. Macrì, M. De Bardi, M. Picozza, I. Salvatori, A. Renzini, S. Valente, A. Mai, V. Moresi, P.L. Puri

**Author notes:** Equal contribution (S. Consalvi, L. Tucciarone).

## Abstract

Pharmacological treatment of Duchenne Muscular Dystrophy (DMD) with histone deacetylase inhibitors (HDACi) is currently being tested in clinical trials. Pre-clinical studies performed in mdx mice - the mouse model of DMD - have shown that HDACi promote compensatory muscle regeneration, while inhibiting fibro-adipogenic degeneration, by targeting fibro-adipogenic progenitors (FAPs); however, these beneficial effects are restricted to early stages of disease progression. We show here that FAPs from late stage mdx mice exhibit epigenetic and transcriptional features of senescence that could not be fully reversed by HDACi. In particular, genome-wide increase in H3K9/14 acetylation at gene promoters of Senescence Associated Secretory Phenotype (SASP) genes was associated with their upregulation in late stage mdx FAPs. Treatment with the HDACi Trichostatin A (TSA) could inhibit SASP gene activation in FAPs, by decreasing H3K9/14 acetylation. Conversely, combinatorial decrease of H3K27 and/or H3K9/14 acetylation at promoters of genes required for cycle activation and progression was associated with their downregulation in FAPs from late stage mdx mice. However, these epigenetic and transcriptional alterations could not be reversed by TSA, due to a general resistance exhibited by FAPs from late stage mdx mice to HDACi-induced H3K9/14 hyperacetylation. Overall, this data reveal that disease-associated features of senescence develop in FAPs of DMD muscle through epigenetically distinct and pharmacologically dissociable events, and suggests that HDACi might at least retain anti- fibrotic and inflammatory activity at late stages of DMD, by repressing FAP-derived SASP.

## Introduction

Duchenne Muscular Dystrophy (DMD) is a fatal genetic disease caused by lack of dystrophin (dys) expression^1,2^. Genetic correction by restoration of dys expression with gene therapy approaches^3–5^ is predicted to recover the biochemical and functional integrity of the dystrophin-associated protein complex (DAPC)^6^, and thereby protect myofiber sarcolemma stability post-contraction^7^. However, a number of “secondary” pathogenic events caused by dys deficiency can contribute to DMD progression^8–13^ and might persist even after gene therapy. Targeting these DMD-associated “secondary” events might therefore be necessary to achieve complete and long-lasting therapeutic recovery in DMD patients.

Among the “secondary” events caused by dys deficiency, pathogenic activation of specific sub-populations of muscle resident cells is emerging as key event in DMD progression^14,15^. Recent works have lended further support to the activation of “secondary” pathogenic responses in cell types that do not express dys, by showing alterations of the transcriptional profiles in various muscle-resident cell types from mdx muscles, in addition to myonuclei and MuSCs^16–20^.

A number of pharmacological strategies have been proposed to target the pathogenic activation of muscle-resident cell types in DMD, including the current standard treatment with steroids^21^. Treatment with HDACi is emerging as novel pharmacological intervention in DMD^22^. The therapeutic potential of HDACi for DMD has been shown by multiple lines of evidence, including preclinical^23,24^ and early clinical studies^25^, and is currently under evaluation in clinical trials with DMD boys^26^. Studies in mdx mice - the DMD murine model – have shown that the beneficial effects of HDACi are restricted to the early stages of disease progression^27,28^. This loss of beneficial effects observed in DMD mouse models at late stages of disease suggests that development of disease-associated resistance might limit the efficacy of HDACi in late stage DMD patients. Among muscle-resident cells, fibro-adipogenic progenitors (FAPs) have been implicated as central cellular effectors of DMD progression and key targets of the beneficial effects of HDACi in mdx mice^27,28^. FAPs support muscle-stem cell mediated repair in acutely injured muscles, but turn into cellular effectors of fibrotic and adipogenic degeneration of muscles exposed to conditions of chronic damage, such as DMD and other neuromuscular disorders^16,27–35^. We have previously shown that in mdx mice at early stages of disease, FAPs can promote muscle stem cell (MuSC)-mediated compensatory regeneration and are susceptible to HDACi-mediated enhancement of their pro-regenerative activity and inhibition of their fibro-adipogenic potential^27,28^. Moreover, recent studies have revealed that exposure to HDACi promotes the formation and release of pro-regenerative and anti-fibrotic extra-cellular vesicles (EVs) from FAPs of DMD muscles at early stages of disease^36^. The progressive loss of pro-regenerative potential and response to HDACi in FAPs from late stages mdx mice^27^ suggests that proportional changes in HDAC activity and related histone modifications occur in these cells during DMD progression. However, it remains currently unknown whether HDAC activity is altered in FAPs of dystrophic muscles and can generate aberrant profiles of histone acetylation and gene expression; likewise, it is unknown whether these alterations could be effectively restored by HDACi at progressive stages of disease progression and what is their impact on FAP biology.

## Results

We measured class I and class II HDAC activity from lysates of FAPs isolated from hind limb muscles of mdx mice at either early (1.5 month old) or late (12 month old) stages of disease – hereinafter also referred to as “young mdx FAPs” or “old mdx FAPs”, respectively. The enzymatic activity of all HDAC classes – HDAC I, IIa and I/IIb – was increased about two folds in mdx old FAPs, as compared to young mdx FAPs (Fig. 1A). A 15 day treatment with the pan HDACi Trichostatin A (TSA), which we have previously reported to exert histological and functional beneficial effects in young mdx mice^23^, could reduce class I HDAC enzymatic activity with a comparable efficacy in mdx FAPs from either stage (Fig. 1A, left panel). By contrast, class IIa HDAC activity was drastically inhibited (about 10 fold reduction) by TSA in young mdx FAPs, whereas it was only reduced by half in old mdx mice (Fig. 1A, middle panel). Finally, class I/IIb HDAC activity was moderately inhibited by TSA in young mdx FAPs and minimally affected in old mdx FAPs (Fig. 1A, right panel). These results show that HDAC activity increases in FAPs of mdx muscles along with disease progression, and that exposure to TSA could reduce the enzymatic activity of class I and II HDAC at both stages, although with a greater activity in young mdx FAPs.

**Figure 1.**
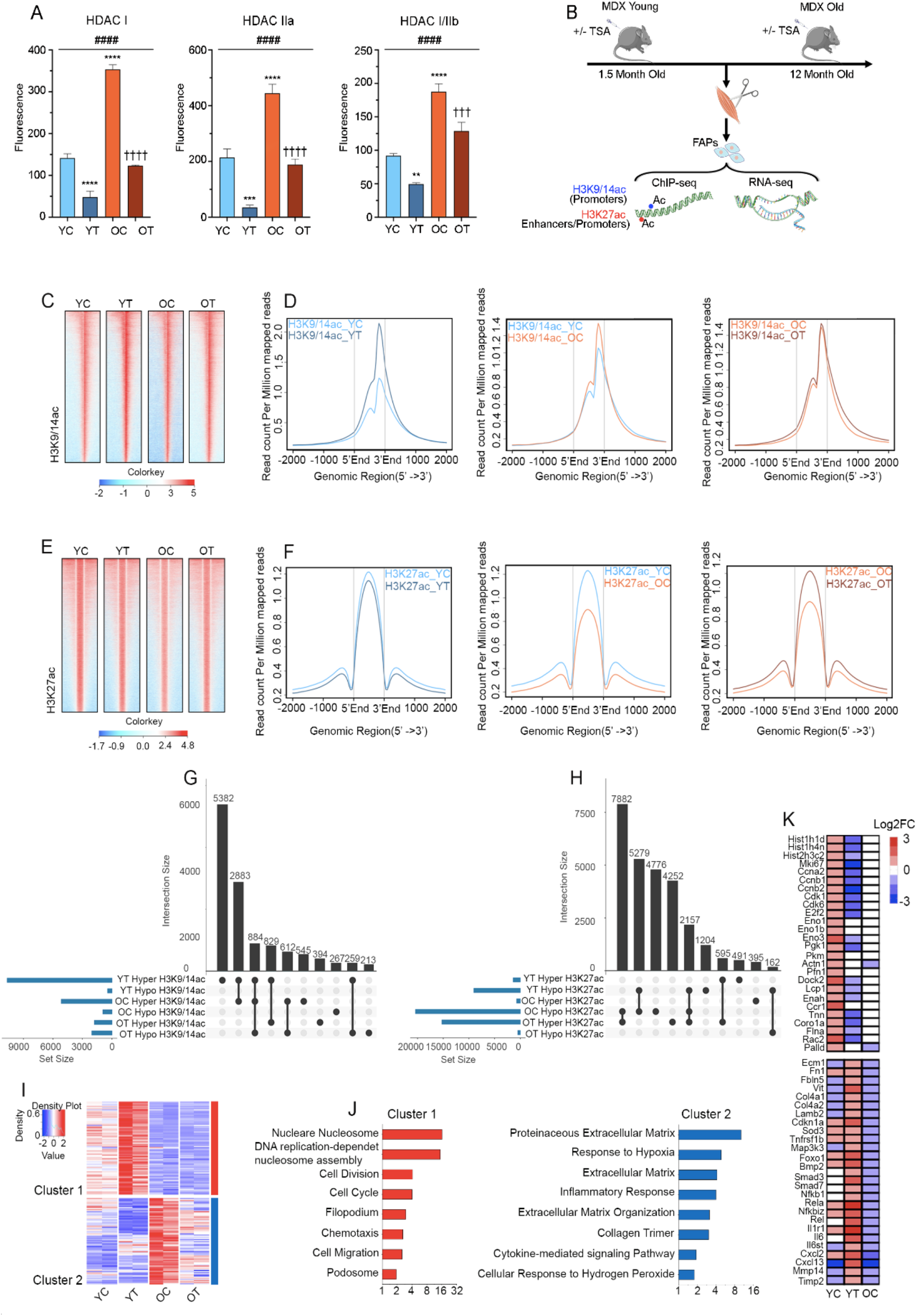
Differential patterns of HDAC activity and gene expression in FAPs during DMD progression and treatment with HDACi. YC: young mdx FAPs; YT: young mdx FAPs *in vivo* treated with TSA for 15 days; OC: old mdx FAPs; OT: old mdx FAPs *in vivo* treated with TSA for 15 days. **A)** Graphs showing the enzymatic activity of class I (left panel), class IIa (middle panel) and class I/IIb (right panel) HDACs performed in FAPs isolated from young and old mdx mice treated or not with TSA. Data are represented as average ± SEM (n=4). ** p ≤ 0.01, **** p ≤ 0.0001, against YC by t-test; ^††††^ p ≤ 0.0001 against OC by t-test; ^####^ p<0.0001 by one way ANOVA. **B)** Cartoon illustrating the experimental strategy. Mdx mice at 1.5 months (young) and 12 months (old) of age were treated with TSA or its vehicle of control for 15 days. At the end of the treatment FAPs were isolated by FACS from hindlimb muscles to perform H3K9/14ac and H3K27ac ChIP-seq and RNA-seq. **C)** Heatmap for H3K9/14ac ChIP-seq signal in the experimental conditions described in B). **D)** NGS plot showing H3K9/14ac comparative patterns in the experimental conditions described in B). **E)** Heatmap for H3K27ac ChIP-seq signal in the experimental conditions described in B). **F)** NGS plot showing H3K27ac comparative patterns in the experimental conditions described in B). **G)** Upset graph showing the intersection size (in black) between the differentially acetylated loci for H3K9/14ac (in blue) in the experimental conditions described in B). The top 10 intersections are shown. **H)** Upset graph showing the intersection size (in black) between the differentially acetylated loci for H3K27ac (in blue) in the experimental conditions described in B). The top 10 intersections are shown. **I)** Heatmap showing 2 clusters of DE genes identified across all the experimental conditions described in B). Gene expression is represented as z-score calculated across the rows. **J)** Gene Ontology performed on cluster 1 (left panel) and cluster 2 (right panel) genes. **K)** Heatmap showing the differential expression levels by log2 fold change for representative genes of cluster 1 (top panel) and cluster 2 (bottom panel).

As HDAC activity controls histone acetylation, we next sought to determine whether the different levels of HDAC activity detected in FAPs from muscles of mdx mice at different stages of disease could generate different profiles of genome-wide distribution of histone 3 (H3) acetylation at lysines 9/14 (H3K9/14ac) and lysine 27 (H3K27ac) - two major histone modification associated with a chromatin conformation permissive for gene expression^37,38^. We also investigated the effect of HDACi on these histone acetylation patterns, by exposing mdx mice to TSA, as described above. In parallel, we performed RNAseq analysis in order to monitor the transcriptional output from FAPs in the same experimental conditions. Figure 1B illustrates the experimental strategy. ChIP-seq experiments with anti-H3K9/14ac and H3K27ac antibodies revealed distinct profiles of histone acetylation in FAPs isolated from hind limb muscles of young (1.5 month) or old (12 month) mdx mice, either untreated or treated with TSA for 15 days (Fig. 1C-F). Global analysis of the cumulative genomic distribution of ChIP-seq peak signals for these histone modifications showed that H3K9/14ac was largely biased toward gene promoters, while H3K27ac signal was distributed between gene promoters (one half) and intronic or distal intergenic elements that typically harbor enhancers (Fig. S1). This signal is consistent with the well established enrichment of H3K27ac at active enhancers and promoters^37–39^. Slightly increased genome-wide levels of H3K9/14ac were observed at gene promoters of old mdx FAPs, as compared to young mdx FAPs (Fig. 1C and 1D, middle panel). Interestingly, TSA treatment increased H3K9/14ac signal at gene promoters in young mdx FAPs (Fig. 1C and 1D, left panel), while did not significantly alter the global H3K9/14ac levels at gene promoters in old mdx FAPs (Fig. 1C and D, right panel). Conversely, a dramatic loss of H3K27 acetylation at both gene promoters and outside was observed in old mdx FAPs, as compared to their younger counterpart (Fig. 1E and 1F, middle panel). TSA treatment could recover H3K27 acetylation in old mdx FAPs (Fig. 1E and 1F, right panel) to levels comparable to those of young mdx FAPs (Fig. 1F, compare middle and right panels). By contrast, TSA decreased H3K27ac signal in young mdx FAPs inside and outside gene promoters (Fig. 1E and 1F, left panel).

Combinatorial intersection of differentially acetylated loci in FAPs across all experimental conditions for both histone modifications shows that the large majority of the H3K9/14ac peaks detected was induced by TSA in young mdx mice, with more than half of them specific for this condition (Fig. 1G, left panel). Most of the remainder H3K9/14ac peaks induced by TSA in young mdx FAPs coincided with H3K9/14 hyperacetylated loci in old mdx FAPs, with a subset of them also coinciding with H3K9/14 hypoacetylated loci induced by TSA in old mdx FAPs (Fig. 1G). This pattern of combinatorial intersections identifies a putative common subset of gene promoters regulated by H3K9/14 acetylation in both young and old mdx FAPs, whereby common hyperacetylated loci detected in TSA-treated young mdx FAPs and untreated old mdx FAPs coincided with loci hypoacetylated in response to TSA treatment in old mdx FAPs. This specific combination suggests that TSA might differentially affect the H3K9/14ac status of a common subset of gene promoters in FAPs throughout DMD progression. Interestingly, another subset of H3K9/14 hyperacetylated loci uniquely detected in old mdx FAPs coincided with hypoacetylated loci in TSA-treated old mdx FAPs (Fig. 1G), further indicating that reversal of H3K9/14 hyperacetylation at gene promoters is a dominant, paradoxical effect of HDACi in old mdx FAPs. Indeed, TSA-mediated H3K9/14 hyperacetylation in old mdx FAPs was a rare event and mostly occurred at loci that were also hyperacetylated by TSA in young mdx FAPs (Fig. 1G). These data suggest that along with the disease progression in mdx mice, FAPs might develop resistance to HDACi-induced H3K9/14 hyperacetylation, while becoming vulnerable to HDAC-mediated reduction of H3K9/14ac signal at hyperacetylated loci in old mdx FAPs. Conversely, the most dominant combinatorial patterns of H3K27ac included the hypoacetylation at gene loci in old mdx FAPs, with about half of these loci in which H3K27 hyperacetylation was recovered by TSA (Fig. 1H). The other half included loci that were also hypoacetylated in young mdx FAPs treated with TSA or loci uniquely detected in old mdx FAPs, in which H3K27ac signal was not recovered by TSA (Fig. 1H). This pattern shows that the reduction of H3K27ac in old mdx FAPs can be recovered by TSA at certain loci, but not at others, thereby indicating that old mdx FAPs develop partial resistance to HDACi-mediated H3K27 hyperacetylation. Interestingly, in young mdx FAPs TSA could only reduce H3K27ac, both at unique loci and at loci that were also hypoacetylated in old mdx FAPs (Fig. 1H).

The alterations of the genome-wide histone acetylation profiles detected in FAPs of mdx mice at different stages of disease and in response to TSA predict that consensual alterations in gene expression profiles could also occur in mdx FAPs in the same experimental conditions. We therefore performed RNAseq to analyze the gene expression profile of FAPs isolated from the same experimental conditions described above (illustrated in Fig. 1B). Differentially expressed (DE) genes between young and old mdx FAPs were almost equally distributed between up- or down-regulated genes (Fig. S2A, middle panel; Fig. S2B). Likewise, TSA induced a similar number of up- and down-regulated DE genes in young mdx FAPs (Fig. S2A, left panel; Fig. S2B). In contrast, TSA preferentially dowregulated gene expression in old mdx FAPs (Fig. S2A, right panel; Fig. S2B). Combinatorial intersection of DE genes in FAPs across all experimental conditions revealed that the majority of them were genes either uniquely up-regulated or downregulated in old mdx FAPs, as compared to their young counterpar<u>t</u>; however, while the expression of a proportion of genes upregulated in old mdx FAPs was recovered by TSA-mediated repression, the expression of very few genes that were downregulated in old mdx FAPs was recovered by TSA-mediated activation (Fig. S2C). Overall, TSA-modulated genes in young and old FAPs did not show any relevant overlap, suggesting that HDACi modulate different patterns of gene expression in dystrophic FAPs at different stages of disease, as also predicted by their acetylation profiles. Heatmap of top DE genes across all experimental conditions revealed specific patterns of gene expression that could discriminate 2 major clusters of DE genes during FAP transition from an early to a late stage of disease progression in mdx mice, as well as their differential response to TSA (Fig. S3 and Fig. 1I). Gene ontology analysis identified specific biological processes for each of these clusters of DE genes. Cluster 1 included a subset of genes whose expression was induced by TSA in young mdx FAPs, but was repressed, and was not recovered by TSA, in old mdx FAPs (Fig. S3 and Fig. 1I). Gene ontology assigned these genes to processes related to activation of cell proliferation and migration (Fig. 1J). These genes encode cell cycle activators, such as cyclins and cyclin-dependent kinases, histone variants, components of the cytoskeleton as well as activators of glycolysis (Fig. S3 and Fig. 1K). Cluster 2 included a subset of genes induced in old mdx FAPs, as compared to young mdx FAPs; however, TSA treatment could downregulate these genes and restore their expression to levels comparable to those of young mdx FAPs (Fig. S3 and Fig. 1I). Gene ontology analysis indicates that this cluster was enriched in genes implicated in regulation of extracellular matrix (ECM), response to hypoxia and inflammation, as well as cytokine-mediated signaling pathways (Fig. 1J). Indeed, these genes encode a number of proteins implicated in ECM remodeling, several ligands and receptors for activation of intracellular signaling as well as downstream nuclear transcription factors (Fig. S3 and Fig. 1K).

We next performed an integrated analysis of ChIPseq and RNAseq datasets generated across all experiemental conditions to identify specific patterns of histone acetylation and gene expression that could discriminate young from old mdx FAPs and their different ability to respond to TSA. Specific patterns of histone acetylation were associated to the expression levels of the nearest gene (−1500/+500 bp distance from the TSS). This analysis revealed two major trends, one consisting of genes that were upregulated in old mdx FAPs and marked by H3K9/14 hyperacetylation at their promoters, another consisting of genes that were downregulated in old mdx FAPs and marked by H3K27 hypoacetylation (Fig. S4A). Gene ontology revealed that upregulated genes marked by H3K9/14 hyperacetylation were enriched in genes belonging to cluster 2 (Fig. S3 and Fig. 1I-K) and implicated in ECM remodeling (e.g. TGFbeta signaling) and cytokine-mediated signaling pathways (e.g. TNFalpha or NFkB signaling) (Fig. S4B). Figure 2A shows the association between increased H3K9/14ac at promoters of representative genes that were upregulated in old mdx FAPs, such as the components of pro-fibrotic TGFbeta signaling, Smad3, Tgfb2 and Tgfbi (Transforming Growth Factor Beta Induced), (Fig. 2A). The upregulation of these representative genes and the increased levels of H3K9/14ac at their promoters in old mdx FAPs, as compared to young mdx FAPs, was validated by independent qPCR (Fig. 2B) and ChIP-qPCR (Fig. 2C) analyses, respectively. Of note, GSEA enrichment plot for genes upregulated in old mdx FAPs and marked by promoter H3K9/14 hyperacetylation revealed their identity as Senescence Associated Secretory Phenotype (SASP) genes (Fig. 2D)^40^. This suggests that old mdx FAPs might acquire a secretory phenotype similar to senescent cells. Interestingly, genes upregulated in old mdx FAPs and marked by promoter H3K9/14 hyperacetylation were also enriched in aging process and negative regulation of cell cycle (Fig. S4B), further suggesting that FAPs at late stages of DMD progression might adopt additional features of cellular senescence, such as cell cycle arrest. Consistenlty, genes downregulated in old mdx FAPs and marked by H3K27 hypoacetylation showed enrichment for biological processes related to activation of cell cycle progression, DNA replication and mitosis (Fig. S4C). Figure 2E shows the association between promoter H3K27 hypoacetylation and downregulation of representative genes in old mdx FAPs, such as the cell cycle activators E2F1, Cdk4 and Check2 (Fig. 2E). The downregulation of these genes and the decreased levels of H3K27ac at their promoters in old mdx FAPs, as compared to young mdx FAPs, were independently validated by qPCR (Fig. 2F) and ChIP-qPCR (Fig. 2G) analyses, respectively. Accordingly, flow cytometry analysis revealed a drastic decrease of cells progressing through S/G2 phases in old mdx FAPs, as compared to young mdx FAPs (Fig. 2H). Likewise, old mdx FAPs exhibited decreased number of apoptotic cells (Fig. 2I), consistent with the decreased susceptibility to apoptosis of cells upon cell cycle withdrawal. However, old mdx FAPs did not exhibit two conventional markers of cellular senescence, such as spontaneous expression beta-galactosidase (Fig. 2J) and upregulation of the cyclin-dependent kinase inhibitors (cdki) p16, although the cdki p21 was increased (see Fig 1K e Fig. S3). Thus, although FAPs exhibit partial features of senescence, they do not fullfill all essential criteria of conventional cellular senescence.

**Figure 2:**
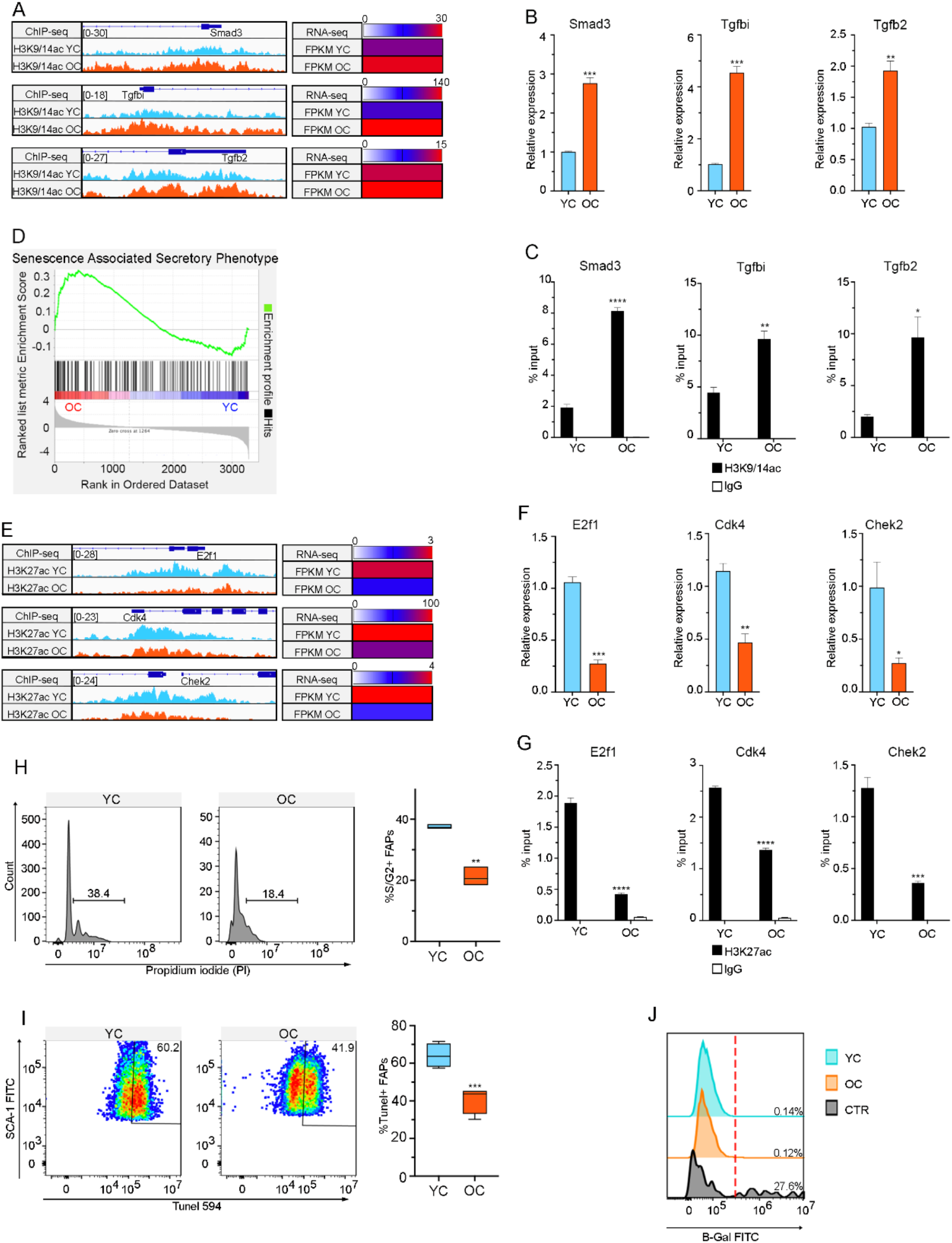
Comparative analysis of FAPs from young and old mdx mice reveals features of cellular senescence during disease progression. YC: young mdx FAPs; OC: old mdx FAPs. **A)** Acetylation tracks for H3K9/14ac ChIP-seq (on the left) and corresponding FPKM values by RNA-seq (on the right) for representative genes in young and old mdx FAPs; **B)** Graphs showing RNA levels for the representative genes showed in A) measured by qPCR; **C)** Graphs showing the H3K9/14 acetylation levels for the representative genes showed in A) measured by ChIP-qPCR. **D)** GSEA enrichment plot of young and old mdx FAPs gene expression profile against the gene list of the Senescence Associated Secretory Phenotype. **E)** Acetylation tracks for H3K27ac ChIP-seq (on the right) and corresponding FPKM values by RNA-seq (on the left) for representative genes in young and old mdx FAPs; **F)** Graphs showing RNA levels for the representative genes showed in E) measured by qPCR; **G)** Graphs showing the H3K27 acetylation levels for the representative genes showed in E) measured by ChIP-qPCR. **H)** Flow cytometry analysis of Propidium Iodide (PI) incorporation to assess the percentage of young and old mdx FAPs in S/G2 phases. Representative plots of PI fluorescence distribution (left panel) and box plot showing the percentage of young and old mdx FAPs in S/G2 (right panel) are shown. **I)** Flow cytometry-based Tunel assay to monitor apoptosis in young and old mdx FAPs. Representative dots of Tunel-594 fluorescence (left panel) and box plot showing the percentage of Tunel-594^+^ FAPs (right panel) are shown. **J)** Flow cytometry analysis of β-Gal staining in young and old mdx FAPs compared to late passage IMR90 senescent cells (CTR). Data are represented as average ± SEM (n=3 for B, C, F and G; n=4 for H-J). *p<0.05, **p<0.01, ***p<0.001, ****p<0.0001 by t-test.

Cellular senescence is traditionally considered irreversible, once established^41^, however recent evidence demonstrates that individual biological features of senescence could be reversible^42,43^. Thus, we sought to evaluate whether epigenetic and transcriptional patterns exhibited by old mdx FAPs were reversed, at least partly, by TSA. In this respect, HDACi have been reported to either promote or counter cellular senescence, depending on the cell type and experimental context^44^. We first evaluated whether TSA could reverse the upregulation of SASP genes in old mdx FAPs. Indeed, the RNAseq expression patterns of cluster 2, which is enriched in SASP genes, shows a consensual downregulation of gene expression levels in FAPs isolated from old mdx mice treated with TSA (Fig. 1 I-K; Fig S3). TSA-mediated downregulation of SASP genes invariably coincided with reduction of H3K9/14ac at their promoters, as shown by representative genes implicated in a variety of SASP-related biological processes, including fibrosis (e.g. TGFb-Smad sgnaling), inflammation (NFkB and p38 signaling) and other components of cytokine- or growth factor-activated pathways, such as IL6, FGF and BMP signaling (Fig 3A; Fig. S5). The downregulation of these genes and the decreased levels of H3K9/14ac at their promoters in response to TSA were independently validated by qPCR (Fig. 3B) and ChIP-qPCR (Fig. 3C) analyses, respectively. Consistently, immunofluorescence analysis of muscle sections show that FAPs (identified as CD90 expressing interstitial cells) from old mdx mice exhibited the activation of NFkB cascade (as measured by nuclear accumulation of the phosphorylated active form of p65 sub-unit) (Fig. 3D-E), p38 pathway (as measured by nuclear accumulation of phosphorylated-p38 alpha kinase) (Fig. 3F-G) and TGFb signaling (as measured by nuclear accumulation of phospho-Smad2/3) (Fig. 3H-I). Treatment with TSA invariably reduced the activation of all these signaling pathways (Fig. 3D-I).

**Figure 3.**
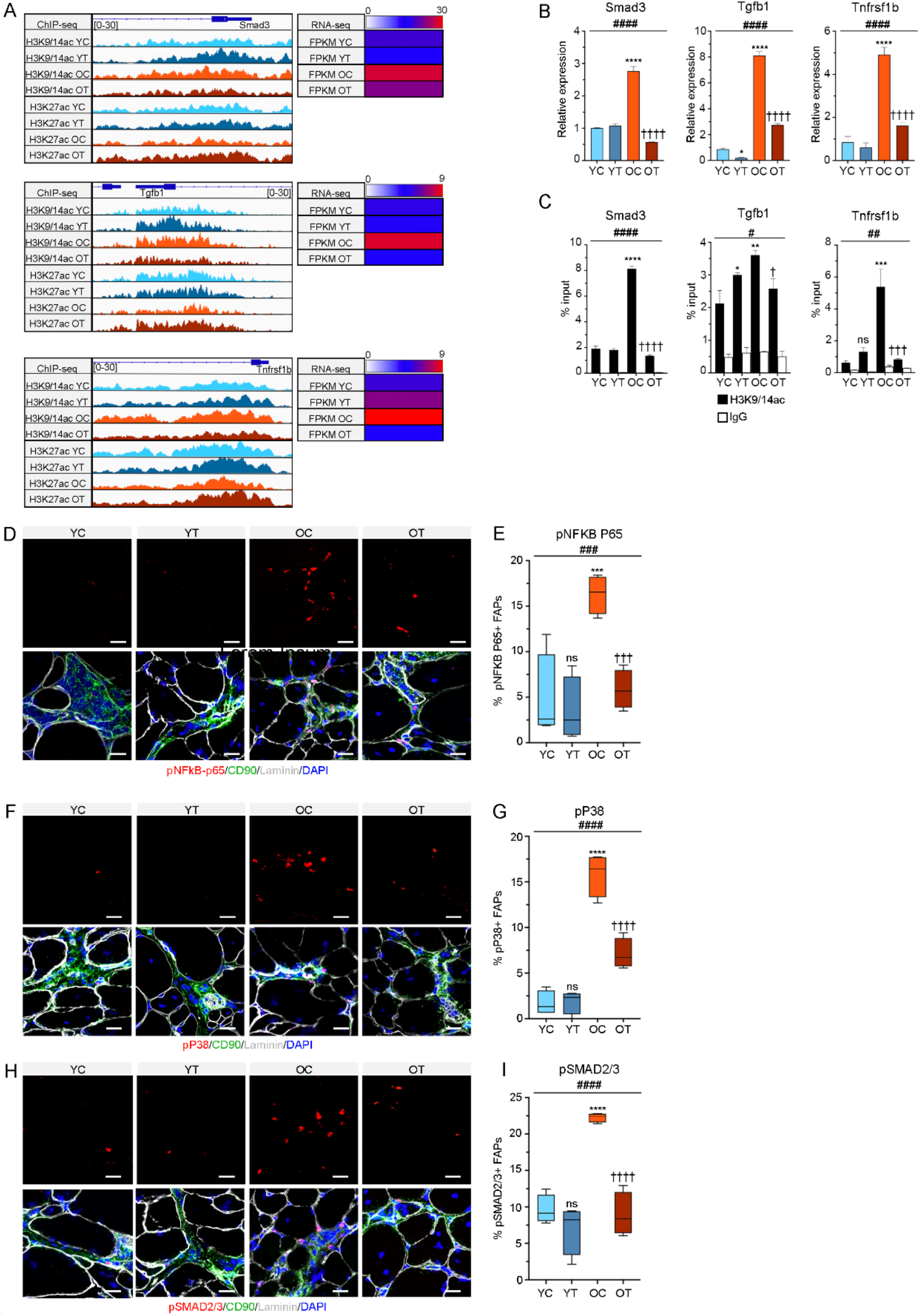
HDACi reduce SASP activation in old mdx FAPs. YC: young mdx FAPs; YT: young mdx FAPs *in vivo* treated with TSA for 15 days; OC: old mdx FAPs; OT: old mdx FAPs *in vivo* treated with TSA for 15 days. **A)** Acetylation tracks for H3K9/14ac and H3K27ac ChIP-seq (on the left) and corresponding FPKM values by RNA-seq (on the right) for representative genes in young and old mdx FAPs treated or not with TSA; **B)** Graphs showing RNA levels for the representative genes showed in A) measured by qPCR; **C)** Graphs showing the H3K9/14 acetylation levels for the representative genes showed in A) measured by ChIP-qPCR. **D)** Representative images of the phospho-NFkB staining in FAPs (pNFkB-p65 in red/CD90 in green/Laminin in grey/DAPI in blue) on tibialis anterior muscle transversal section of young and old mdx mice treated or not with TSA; Scale bar = 25 μm; **E)** Box plot showing the percentage of pNFkB-p65 positive FAPs in the staining shown in D). **F)** Representative images of the phospho-p38 staining in FAPs (pP38 in red/CD90 in green/Laminin in grey/DAPI in blue) on tibialis anterior muscle transversal sections of young and old mdx mice treated or not with TSA. Scale bar = 25 μm; **G)** Box plot showing the percentage of pP38 positive FAPs in the staining shown in F). **H)** Representative images of the phospho-SMAD2/3 staining in FAPs (pSMAD2/3 in red/CD90 in green/Laminin in grey/DAPI in blue) on tibialis anterior muscle transversal sections of young and old mdx mice treated or not with TSA. Scale bar = 25 μm; **I)** Box plot showing the percentage of pSMAD2/3 positive FAPs in the staining shown in H). Data are represented as average ± SEM (n=3); *p<0.05, **p<0.01, ***p<0.001, ****p<0.0001 against young mdx FAPs by t-test; ^†^p<0.05, ^†††^p<0.001, ^††††^p<0.0001 against old mdx FAPs by t-test; ^#^p<0.05, ^##^p<0.01, ^####^ p<0.0001 by one way ANOVA.

Conversely, TSA could not resume the expression of genes downregulated in old mdx FAPs and represented in cluster 1, which is enriched in genes implicated in the activation of cell cycle progression (e.g. cyclins, cdks, histones, E2F family members) as well as glycolysis (Fig. 1 I-K; Fig S3). The common epigenetic features that accompanied the downregulation of these genes during the transition of FAPs from young to old mdx mice was the reduced promoter H3K9/14ac, often in combination with reduced H3K27ac (Fig. 4A; Fig. S6). While TSA could increase H3K9/14 promoter acetylation and expression of cell cycle genes in young mdx FAPs, it failed to recover promoter H3K9/14 hyperacetylation and resume the expression of these genes in old mdx FAPs (Fig. 4A and Fig. S6), as also measured by qPCR and ChIPqPCR analysis at representative genes, such as Hist1h2ae/Hist1h2bg, Cdk1 and CyclinA1 (encoded by Ccna2) (Fig. 4B and C). Consistently, the percentage of proliferating cells detected by flow cytometry analysis of Ki-67 expression – a nuclear protein expressed throughout the cell cycle progression, except G0 and early G1 phases - was slightly reduced in old mdx FAPs, as compared to young mdx FAPs (Fig. 4D and E). However, while TSA could double the percentage of Ki-67 positive FAPs in young mdx mice, old mdx FAPs did not resume the cell cycle in response to TSA treatment (Fig. 4D and E). Accordingly, the total number of FAPs was decreased in old mdx mice, as compared to their young counterpart; however, TSA treatment could not increase FAP number at either stage (Fig. 4F). The lack of increase in FAP number in TSA-treated young mice is apparently in contrast with the increased number of proliferating FAPs detected in TSA-treated young mdx mice. We argue that this discrepancy could be accounted by the increased length of G1-S phase progression observed in TSA-treated young mdx FAPs, due to the activation of a G1/S phase checkpoint by TSA reported by others in several cell types^45^ and confirmed by our RNAseq analysis (see Fig. 5B). These data indicate that old mdx FAPs are withdrawn from the cell cycle, a biological feature that is typically associated with an impairment in migratory ability. We therefore analysed the effect of TSA on FAP migration in young vs old mdx mice, by a cell migration assay in vivo, using FAPs isolated from young or old mdx mice, treated or not with TSA for 15 days. Immediately after isolation, FAPs were labelled with PKH67-488 dye and transplanted into the proximal part of the gastrocnemius muscle of mdx young mice. 5 days post-injection the number of PKH67-488 positive cells detected in the proximal and distal sections from the injection site were counted by flow cytometry analysis, as readout of their migratory ability. Figure 4G-H shows that only FAPs isolated from TSA-treated young mdx mice were able to migrate from the proximal injection site to the distal section of gastrocnemious muscles (Fig. 4G-H), indicating that old mdx FAPs are refractory to TSA-induced migration. Finally, the observation that the expression of several glycolytic genes, including Eno3 (the muscle-specific isoform of beta-enolase), was repressed in old mdx FAPs and could not be resumed by TSA, which otherwise upregulates these genes in young mdx FAPs (Fig. S3), prompted an interest to analyse the ability of FAPs to activate glycolysis in our experimental conditions. Activation of glycolysis could be observed only in FAPs isolated from young mdx mice treated with TSA (Fig. 4I). Given the functional interdependence between glycolysis and activation of cell cycle during stem cell activation, and because cell cycle progression and DNA replication favor nuclear reprograming^47^, it is likely that the resistance of old mdx to resume expression of cell cycle and glycolytic genes prevents full epigenetic reprogramming by HDACi.

**Figure 4.**
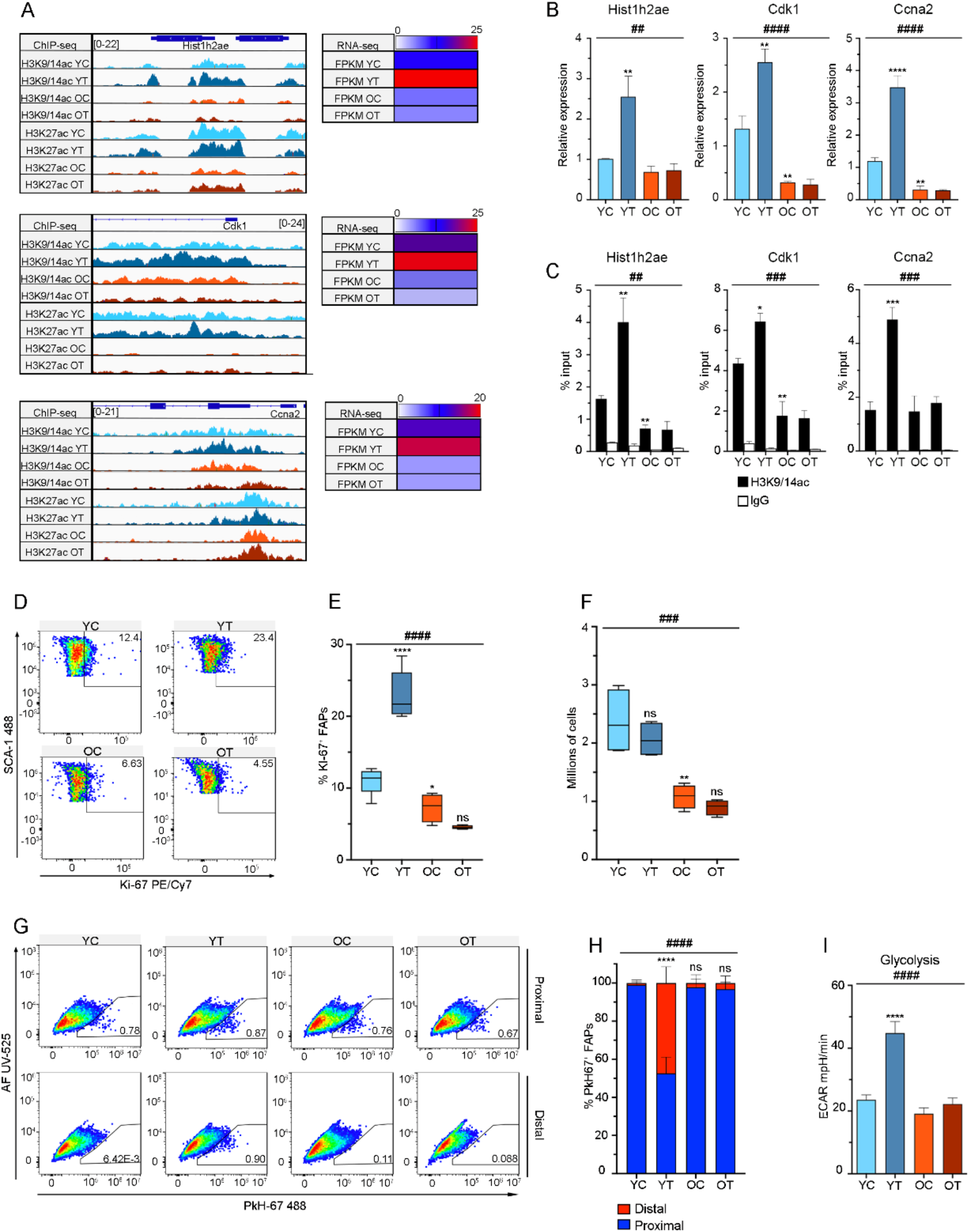
FAPs from old mdx mice develop resistance to HDACi-mediated activation of cell cycle. YC: young mdx FAPs; YT: young mdx FAPs *in vivo* treated with TSA for 15 days; OC: old mdx FAPs; OT: old mdx FAPs *in vivo* treated with TSA for 15 days. **A)** Acetylation tracks for H3K9/14ac and H3K27ac ChIP-seq (on the left) and corresponding FPKM values by RNA-seq (on the right) for representative genes in young and old mdx FAPs treated or not with TSA; Graphs showing RNA levels for the representative genes showed in A) measured by qPCR; Graphs showing the H3K9/14 acetylation levels for the representative genes showed in A) measured by ChIP-qPCR. **D)** Representative dot-plots of the flow cytometry analysis of KI-67 PE-Cy7 to monitor proliferating FAPs in young and old mdx FAPs treated or not with TSA; **E)** Box plot showing the percentage of KI-67^+^ FAPs analysed in D). **F)** Box plot showing the total number of FAPs in the same experimental conditions described in D). **G)** Representative dot-plots showing the flow cytometry analysis of FAP migration assay. Young and old mdx FAPs with or without TSA treatment were labelled with PKH67-488 and analysed in the proximal (top panel) and distal (bottom panel) sections from the injection site in young mdx mice. **H)** Stacked bar chart showing the relative percentage of PKH67-488^+^ FAPs detected in the proximal (blue) and the distal (red) sections for the same experimental conditions described in G). **I)** Histogram showing the level of glycolysis measured by Seahorse as extracellular acidification rate (ECAR) after glucose administration to young and old mdx FAPs treated or not with TSA. Data are represented as average ± SEM (n=3 for B and C; n=4 for D-I). *p<0.05, **p<0.01, ***p<0.001, ****p<0.0001 against young mdx FAPs by t-test; ^##^p<0.01, ^###^p<0.001, ####p<0.0001 by one way ANOVA.

**Figure 5.**
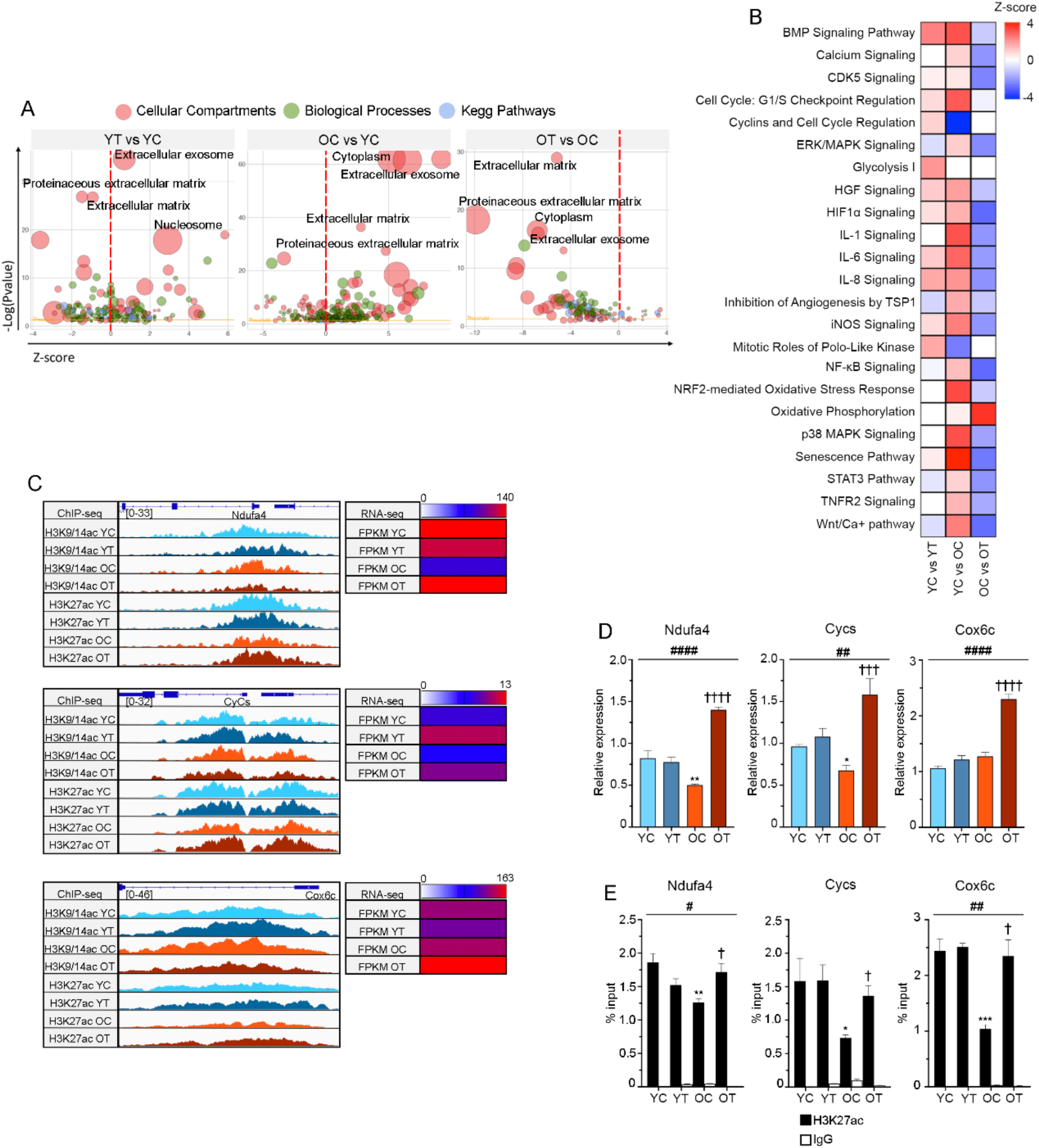
HDACi promotes oxidative phosphorylation in old mdx FAPs. YC: young mdx FAPs; YT: young mdx FAPs *in vivo* treated with TSA for 15 days; OC: old mdx FAPs; OT: old mdx FAPs *in vivo* treated with TSA for 15 days (OT). **A)** Bubble plots showing the Gene Ontology analysis of RNA-seq performed by DAVID in young mdx FAPs treated or not with TSA (left panel), young and old untreated mdx FAPs (middle panel) and old mdx FAPs treated or not with TSA (right panel). The size of the bubbles accounts for the number of genes found to be involved in the GO terms. **B)** Heatmap showing a selection of canonical pathways predicted by IPA in a comparative analysis of young and old mdx FAPs treated or not with TSA. **C)** Acetylation tracks for H3K9/14ac and H3K27ac ChIP-seq (on the left) and corresponding FPKM values by RNA-seq (on the right) for representative genes in young and old mdx FAPs treated or not with TSA; **D)** Graphs showing RNA levels for the representative genes showed in C) measured by qPCR; **E)** Graphs showing the H3K27 acetylation levels for the representative genes showed in C) measured by ChIP-qPCR. Data are represented as average ± SEM (n=3). *p<0.05, **p<0.01, ***p<0.001, against young mdx FAPs by t-test; ^†^p<0.001, ^†††^p<0.001, ^††††^p<0.0001 against old mdx FAPs by t-test; #p<0.05, ^##^p<0.01, ^####^p<0.0001 via one way ANOVA.

Overall, these data reveal a general trend of loss of response to HDACi-mediated activation of gene expression in old mdx FAPs, despite these cells retained the ability to dowregulate gene expression (e.g. SASP genes) in response to HDACi. This trend was further revealed by a gene ontology analysis of comparative gene expression patterns in FAPs across our experimental conditions, in graphical representation whereby the size of the bubbles accounts for the number of genes found to be involved in the GO term (Fig. 5A). This analysis shows that TSA could both inhibit and promote a variety of cellular processes in young mdx FAPs; however, in old mdx FAPs TSA activity was only inhibitory toward the biological processes induced in FAPs during DMD progression (Fig. 5A). Among them, we noted processes implicated in fibrosis (e.g. remodelling of extracellular matrix) as well as extracellular exosome formation/secretion. IPA analysis also documented the general trend of inhibition of gene expression by TSA in old mdx mice, with the notable exception of a cluster of TSA-induced genes implicated in oxidative phosphorylation (Ox-Phos), TCA cycle, electron transport and mitochondrial biogenesis (Fig. 5B; Fig. S7A and B). The dominant epigenetic feature of these genes was the H3K27 hypoacetylation at their promoters that accompanied their downregulation in old mdx FAPs, as compared to young mdx FAPs (Fig. 5C-E; Fig. S7C). TSA could recover promoter H3K27 hyperacetylation and transcription of these genes in mdx old FAPs (Fig. 5C; Fig. S7C). Independent qPCR and ChIPqPCR analysis of representative genes - Ndufa4, Cycs and Cox6 - confirmed their downregulation (Fig. 5D) and consensual reduction of H3K27ac promoter levels (Fig. 5E) in old mdx FAPs. These gene expression patterns were paralleled by coherent changes in mitochondrial activities, with a general trend of reduction of oxygen consumption rate (basal respiration) and ATP production in old mdx FAPs, which was recovered by TSA, albeit not to the levels observed in TSA-treated young mdx FAPs (Fig. S7D and E). Finally, mitosox staining detected an increased production of superoxide in old mdx FAPs, as biproduct of dysfunctional mitochondrial oxydative phosphorylation, that was reverted by TSA (Fig. S7F and G). Interestingly, these genes were not significantly induced by TSA in young mdx FAPs (Fig. 5C-E; Fig. S7C). This suggests that full metabolic reprogramming of young mdx FAPs is induced by TSA by a mechanism distinct from TSA-mediated upregulation of Ox-Phos genes in old mdx FAPs, which can only promote a slightly increase of mitochondrial oxidative phosphorylation (Fig. S7D-G).

## Discussion

The results shown here revealed that FAPs of mdx mice undergo extensive epigenetic and transcriptional changes during disease progression, leading to two main distinctive biological features – cell cycle arrest and SASP - that discriminate late from early stage FAPs.

Although cell cycle arrest and SASP are well known features of cellular senescence^41^, old mdx FAPs did not exhibit conventional hallmarks of cellular senescence, such as beta galactosidase expression and upregulation of the cyclin-dependent kinase inhibitors (cdki) p16. Indeed, the cell cycle arrest observed in old mdx FAPs appears to be caused by a failure to activate the expression of genes implicated in cell cycle progression and DNA synthesis, rather than by the upregulation of p16, as typically observed in cellular senescence. Moreover, SASP activation in old mdx FAPs entails the upregulation of genes encoding for growth factors, cytokines, and other secreted proteins implicated in the regulation of muscle regeneration, inflammation, and ECM remodeling. Some of these genes were already expressed at low levels in young mdx FAPs and, when induced at moderate levels by HDACi, some of these genes could promote environmental conditions conducive to muscle regeneration – e.g. transient inflammation and ECM changes that favor MuSC migration and proliferation. In old mdx FAPs the simultaneous upregulation and persistent expression of all these genes ultimately leads to fibrosis and inflammation, which negatively impact on MuSC-mediated regeneration. Thus, the activation of SASP observed in old mdx FAPs reflects changes in magnitude of transcription of a large collection of genes implicated in regeneration, inflammation and ECM remodelling. Consistently, similar features of cellular senescence have been independently observed in FAPs of mice exposed to exercise-induced muscle damage and were associated to the ability of FAPs to promote muscle regeneration^48^.

Interestingly, cell cycle arrest and activation of SASP in old mdx FAPs were sustained by opposite patterns of histone acetylation. Cell cycle arrest was associated with global hypoacetylation at promoters of repressed genes implicated in cell cycle progression and DNA synthesis. Activation of SASP was sustained by promoter H3K9/14 hyperacetylation of SASP genes. Furthermore, transcriptional repression of cell cycle genes was part of a trend of genome-wide H3K27 hypoacetylation in old mdx FAPs that coincided with a general increased HDAC activity and could be partly reversed by HDACi at the genome-wide level; however, HDACi could not recover the expression of cell cycle genes in old mdx FAPs. Conversely, activation of SASP genes occurred within a trend of H3K9/14 hyperacetylation that was not reversed by HDACi at genome-wide level; however, HDACi could fully repress the activation of SASP genes, by promoting H3K9/14 hypoacetylation at their promoters. The genome-wide increase in H3K9/14ac observed in old mdx FAPs is in apparent conflict with the increased HDAC activity. Likewise, the reduction of H3K9/14ac observed at promoters of SASP genes in FAPs of late stage mdx mice exposed to TSA appears at odd with the expected ability of HDACi to promote hyperacetylation. We argue that these paradoxical effects are likely accounted by the ability of HDACi to target multiple acetylation-dependent and independent events^49–52^ as well as by the complexity of HDACi activity *in vivo*. Indeed, the final outcome of the systemic exposure of FAPs to HDACi depends on both direct and indirect effects, with the latter likely being generated by signals derived from other cell types that are simultaneously exposed to HDACi. This is particularly relevant for experiments that require long-term exposure to HDACi, as the treatment of mdx mice with TSA. Indeed, parallel RNAseq analysis of macrophages and MuSCs of mdx mice exposed to TSA treatment also show dramatic changes in gene expression (SC, LT and PLP unpublished data). Hence, it is likely that the overall effects of HDACi on genome-wide histone acetylation and transcriptional output in mdx FAPs is determined by modulation of the heterotypic interactions that FAPs establish with muscle-resident cell types. Conceivably, the senescence-associated features observed in late-stages mdx FAPs might result from functional interactions with other cell types implicated in DMD pathogenesis; in turn, FAP-derived SASP can promote the survival or expansion of other cell types that contribute to DMD progression. Therefore, the permanent cell cycle arrest and SASP observed in FAPs at late stages of DMD should be considered as disease-associated features of senescence induced by pathogenic signals released from DMD muscles, rather than conventional features of cell-autonomous activation of cellular senescence. This contention is also supported by the observation that cell cycle arrest and activation of SASP could be already observed in FAPs of mdx mice older than 5 months, but could not be detected in FAPs of aged (older than 2 years) wild type mice^52^. Overall, the ability of HDACi to promote either moderate and reversible activation of specific SASP genes in young mdx FAPs or global repression of the SASP genes in old mdx FAPs appears a major determinant of the different therapeutical effects of HDACi at early vs late stages of DMD progression. In particular, the global repression of SASP genes by HDACi in old mdx FAPs might account for the resistance to the pro-regenerative effects of HDACi observed at late stages of disease, but also indicates that HDACi might retain anti-fibrotic and anti-inflammatory effects at late stages of DMD. This therapeutic “trade off” is due to the general trend of old mdx FAPs toward a resistance to HDACi-mediated activation of gene expression and release of other secretory signals implicated in muscle regeneration (Fig. 5A and B), including release of pro-regenerative EVs^36^.

The differential effects of HDACi observed in young vs old mdx FAPs were associated to their ability to activate cell cycle, glycolysis as well as Ox/Phos metabolism in young, but not old, mdx FAPs. Dysfunctional mitochondrial metabolism has been previously reported in mdx FAPs and correlates with an increased adipogenic potential^53^. Although HDACi could induce the expression of Ox/Phos genes in old mdx FAPs, this effect was not sufficient to activate basal mitochondrial respiration and ATP production at the extent observed in young mdx FAPs exposed to HDACi (Fig. 5 and Fig. S7). Given the intimate link between cell cycle progression, mitochondrial activity, acetyl-coA metabolism and availability of acetyl groups for histone acetylation^47,54^, it is possible HDACi-mediated activation of cell cycle and mitochondrial activities in young mdx FAPs enable histone hyperacetylation at promoters of genes required for pro-regenerative activities. Conversely, reduced availability of mitochondria-derived acetyl-CoA in old mdx FAPs might limit their ability to respond to HDACi with an increased hyperacetylation at gene promoters.

Nonetheless, the HDACi-mediated activation of Ox/Phos genes in old mdx FAPs and repression of SASP genes could be functionally associated, by the suppression of oxidative stress, as recently proposed in models of cellular senescence^44^.

Overall, our data provide evidence that cell cycle arrest and SASP are two senescence-associated biological features that limit the response to HDACi in old mdx FAPs. However, their pharmacological dissociation by HDACi (repression of SASP, without reactivation of cell cycle) suggest that HDACi can exert anti-fibrotic and anti-inflammatory effects also at late stages of DMD progression.

## Materials and Methods

### Animals and in vivo treatments

Mice were bred, handled and maintained according to the standard animal facility procedures and the internal Animal Research Ethical Committee according to the Italian Ministry of Health approved experimental protocols and in agreement with the ethic committee of the Fondazione Santa Lucia (FSL) approved protocols.

C57Bl6 mdx mice were purchased from Jackson Laboratories. C57Bl6 mdx mice at 1.5 and 12 months of age (respectively defined young and old mdx mice) were treated for 15 days with daily intra peritoneal injections of Trichostatin A, TSA (0.6 mg/kg/day; #T8552, Sigma), dissolved in saline solution or in saline alone as vehicle control (CTR).

For the migration assay, FAPs isolated from young or old mdx mice treated or not with TSA were stained with PKH67-488 (#MINI-67, Sigma) immediately after isolation by FACS and injected (20 μl at the concentration of 5000 cells/μl) in the proximal part of the gastrocnemius from the foot-paw of young mdx mice. Mice were sacrificed 5 days post-injection. Gastrocnemius was harvested and cut in half to obtain proximal and distal sections from the injection site. PKH67-488^+^ FAPs were detected by flow cytometry.

### FACS isolation of FAPs

FAPs were isolated from C57Bl6 mdx mice at the end of the treatments immediately after the sacrifice. Hind limb muscles for each mouse were minced and put into a 15 ml tube containing 4 ml of digest solution in HBSS (#24020-091,GIBCO) containing 2 mg/ml Collagenase A (#10103586001, Roche), 2.4 U/ml Dispase II (#04942078001, Roche), 10 ng/ml DNase I (#11284932001, Roche) for 90 min at 37°C. Cells were filtered through 100um, 70um and 40um cell strainers (#08-771-19, #08-771-2, #08-771-1, BD Falcon) and resuspended in 0.5 ml of HBSS containing 0.2% w/v BSA and 1% v/v Penicillin–Streptomycin for the staining of cell surface antigens 30 min on ice. The following antibodies were used: CD45-eFluor 450 (1:50, #48-0451-82, eBiosciences), CD31-eFluor 450 (1:50, #48-0311-82, eBioscience), Ter119-eFluor 450 (1:50, #48-5921-82, eBiosciences), Itga7-649 (1:500, #67-0010-01, AbLab) and Sca1-FITC (1:50, 5981-82, eBioscience). Cells were finally washed and resuspended in 1 ml of HBSS containing 0.2% w/v BSA and 1% v/v Penicillin–Streptomycin. FAPs were isolated as Ter119^-^/CD45^-^/CD31^-^/ Itga7^-^/Sca1^+^ cells using a Beckman Coulter MoFlo Legacy high-speed cell sorter.

### Flow cytometry analysis

Hind limb muscles were digested and the cells were stained with antibodies for cell surface antigens as previously described in the FACS protocol. Then cells were fixed in 4% Formaldehyde (30 min, RT) and washed in PBS. For cell cycle analysis, Propidium Iodide (PI) DNA staining was performed (5ul of PI and 2,5ul of RNAse in 500ul of PBS, 30 min at 37°C in the dark). For apoptosis analysis, Click-iT^®^ Plus TUNEL Assay (Alexa Fluor™ 594 dye, C10618, Termofisher) was performed following manufacturer indications. For senescence analysis, CellEvent™ Senescence Green Detection Kit (C10850, Termofisher) was used following manufacturer indications. For cell cycle activation analysis, staining with Ki-67 PE/Cy7 (652426, Biolegend) was used. For migration assay, total FAPs were detected as Sca1-APC/Fire 750-positive cells (1:100, #108145, Biolegend) and injected FAPs were discriminated as PKH67-488^+^ cells. Cell suspensions were acquired using a CytoFLEX LX flow cytometer (Beckman Coulter) and data were analyzed using FlowJo software (BD Biosciences).

### HDAC enzymatic activity assay

HDAC activity was evaluated by using different fluorogenic substrates specific for class I, class IIa or Class I/IIb HDACs^92^. The assay was performed as previously described^93^. Briefly, freshly isolated FAPs were suspended in PBS (pH 7.4) containing 0.5% Triton X-100, 300 mM NaCl and protease/phosphatase inhibitor cocktail (Thermo Fisher Scientific) and sonicated prior to clarification by centrifugation. Protein concentrations were determined using a BCA Protein Assay Kit (Thermo Fisher Scientific). Extracts were diluted into PBS buffer in 100 μl total volumes in 96-well plate (8 μg FAPs protein/well). Substrates were added (5 μl of 1 mM DMSO stock solution), and the plates were returned to the 37°C incubator for 3 hours. Then, developer/stop solution was added (50 μl per well of PBS with 1.5% Triton X-100, 3 μM TSA, and 0.75 mg/ml trypsin), with additional 20’ incubation at 37°C. To detect fluorescent signal Glowmax (Promega) instrument with excitation and emission filters of 360 nm and 460 nm, respectively was used. Background signals from buffer blanks were subtracted.

### MitoSOX assay

Freshly isolated FAPs were plated in culture media (BIO-AMF-2, Biological Industries) at high density (3,000 cells, in 96-well dishes). After 24 hrs FAPs were treated with MitoSOX reagent (Red Mitochondrial Superoxide Indicator, Thermo Fisher, #M36008) and Hoecst 33342 solution (Thermo Fisher, # 62249) following the manufacturer protocols. Images were acquired using Zeiss LSM 800 confocal microscope and quantified using ImageJ software.

### Real-time cell metabolic analysis

Mitochondrial function and glycolysis rate were determined using a Seahorse XF96e Analyzer (Seahorse Bioscience - Agilent, Santa Clara CA, USA). FAPs, 10,000 per well, were plated on cell tak (2 μg per well; Corning^®^) coated Seahorse 96-well utility plate, centrifuged at 1100 rpm for 10 minutes at room temperature and held at 37 ° C in a CO_2_-free incubator for 45 minutes prior to testing.

Mitochondrial function was assessed through a Cell Mito Stress test. Growth medium was replaced with XF test medium (Eagle’s modified Dulbecco’s medium, 0 mM glucose, pH=7.4; Agilent Seahorse) supplemented with 1 mM pyruvate, 10 mM glucose and 2 mM L-glutamine. The test was performed by measuring at first the baseline oxygen consumption rate (OCR), followed by sequential OCR measurements after injection of oligomycin (1 μM), carbonyl cyanide 4-(trifluoromethoxy) phenylhydrazone (1 μM) and Rotenone (0,5 μM) + Antimycin A (0,5 μM) to obtain the key parameters of the mitochondrial function including basal respiration and ATP-linked respiration.

The rate of glycolysis was determined by measuring the rate of extracellular acidification (ECAR). Cells were cultured and pretreated as previously described. The growth medium was replaced with XF test medium (Eagle’s modified Dulbecco’s medium, 0 mM glucose, pH=7.4; Agilent Seahorse) supplemented with L-glutamine (1 mM). ECAR was repeatedly evaluated after the injection of glucose (10 mM), oligomycin (1 μM) and 2D Glucose (50mM) respectively, in each well.

XF96 data were calculated using the algorithm described and used by the Seahorse software package.

### Histology and immunofluorescence

Tibialis anterior muscles were snap frozen in liquid nitrogen-cooled isopentane and then cut transversally with a thickness of 8 μm. For immunofluorescence analysis, cryo-sections were fixed in 4% PFA for 10 min and permeabilized with 0,25% Triton for 15 min at RT. Muscle sections were blocked for 1h with a solution containing 4% BSA (#A7030, Sigma) in PBS and then incubated with primary antibodies O.N. at 4°C. Antibody binding was revealed using secondary antibodies coupled to Alexa Fluor 488, 594, or 647 (Invitrogen). Sections were incubated 5 min with DAPI in PBS for nuclear staining, washed in PBS, and mounted with glycerol 3:1 in PBS. The primary antibodies used for immunofluorescences are: rabbit anti-Laminin (1:400, L9393, Sigma), rabbit anti-Phospho-NFκB p65 (1:1000, 3033, Cell Signalling,), rabbit anti-Phospho-p38 MAPK (1:1000, 4511, Cell Signalling), rabbit anti-Phospho-Smad2 (Ser465/467)/Smad3 (Ser423/425) (1:1000, 8828, Cell Signalling).

### RNA-seq sample preparation

FAPs were isolated by FACS from 6 C57Bl6J mdx male mice for each experimental condition. RNA was collected using Trizol reagent (#T9424, Sigma) and 1 ug (100 ng/ul) were sent in duplicate to IGA (Istituto di Genomica Applicata, Udine) for RNA sequencing using Illumina TruSeq Stranded Total RNA kit Ribo-Zero GOLD on Illumina Hiseq2500 platform.

### RNA-seq validation

Total RNA was extracted from freshly isolated FAPs using Trizol and 0.5–1mg were retro-transcribed using the Taqman reverse transcription kit (Applied Biosystems). Real time quantitative PCR(RT-qPCR) was performed to analyse relative gene expression levels using SYBR Green Master mix (Applied Biosystems) following manufacturer indications. Relative expression values were normalized to the housekeeping gene GAPDH.

Primers sequences are as follow:

CCNA2:
Fwd: GTCCTTGCTTTTGACTTGGC
Rev: ACGGGTCAGCATCTATCAAAC

CDK1:
Fwd: TGCAGGACTACAAGAACACC
Rev: GCCATTTTGCCAGAGATTCG

CDK4:
Fwd: ACAAGTAATGGGACCGTCAAG
Rev: GGGTGTTGCGTATGTAGACTG

CHEK2:
Fwd: CTGAGGACCAAGAACCTGAAG
Rev: CCATCGAAGCAATATTCACAGC

COX6C:
Fwd: TGCGGGTTCATATTGCTGG
Rev: CAGCCTTCCTCATCTCTTCG

CYCS:
Fwd: CCAAATCTCCACGGTCTGTTC
Rev: ATCAGGGTATCCTCTCCCCAG

E2F1:
Fwd: TCTCTTTGACTGTGACTTTGGG
Rev: TCGTGCTATTCCAATGAGGC

GAPDH:
Fwd: CACCATCTTCCAGGAGCGAG
Rev: CCTTCTCCATGGTGGTGAAGAC

HIST1H2AE:
Fwd: CACATCAGCTTTTCCACTTCCA
Rev: GTCCAGACATTGACGCAAGAAG

NDUFA4:
Fwd: TGCGCTTGGCACTGTTTAATC
Rev: AGTCTGGGCCTTCTTTCTTCA

SMAD3:
Fwd: CCGAGAACACTAACTTCCCTG
Rev: CATCTTCACTCAGGTAGCCAG

TGFBI:
Fwd: AACCGACCACAAGAACGAG
Rev: GCTTCATCCTCTCCAGTAACC

TGFB1:
Fwd: CCTGAGTGGCTGTCTTTTGA
Rev: CGTGGAGTTTGTTATCTTTGCTG

TGFB2:
Fwd: TGCTAACTTCTGTGCTGGG
Rev: GCTTCGGGATTTATGGTGTTG

TNFRSF1B
Fwd: ACTCCAAGCATCCTTACATCG
Rev: TTCACCAGTCCTAACATCAGC

### RNA-seq analysis

RNA sequencing analysis was performed mapping more than 20 millions of reads for each sample to the Mus Musculus GRCm38.78 genome using TopHat 2.0.9. Read count was performed with HTSeq-0.6.1p1. Mapped reads were analyzed with R-studio (R version 3.5.2) using DESeq2 to obtain differentially expressed (DE) genes with normalized RPKM, p-value, p-adjusted and log2fold change values. Genes were considered differentially expressed for p-adjusted < 0.1.

DE genes were visualized by Heatmap, MA plots (generated with DESeq2) and Violin plot (generated with Prism 8.0).

Deseq2 differential genes were analysed for overlap between datasets using the Intervene tool with default parameters. The matrix of overlaps was generated in R 3.5.2 and uploaded in the intervene shiny app (https://asntech.shinyapps.io/intervene/) and the Upset graph was generated.

Clusters were manually curated from the total RNA-seq heatmap. For gene ontology DE genes (p-adj<0.1) were uploaded in https://david.ncifcrf.gov and the most relevant GO terms were manually selected. Results were processed in the R package GOplot for the bubble plots. The associated gene expression heatmap was generated by manually selecting the most relevant genes for each biological category from David Gene Ontology. Heatmaps and histograms were generated in Graphpad prism 8.0.

QIAGEN Ingenuity Pathway Analysis (IPA) was performed as comparative analysis of the multiple experimental groups filtering DE genes for p-adj<0.1. A selection of significant canonical pathways (p-value<0.1) was shown as heatmap generated in Graphpad prism 8.0. Gene set enrichment analysis (GSEA) was performed using genes H3K9/14 hyper-acetylated in old versus young mdx FAPs. Weighted statistic was used to reveal significant enrichment of SASP genes. A list of SASP transcripts was developed and compiled from several sources studying senescence^94,95^.

### ChIP-seq sample preparation

FAPs were freshly isolated by FACS from 10 C57Bl6J mdx male mice for each experimental condition. DNA was double-crosslinked to proteins with 37% formaldehyde (Sigma) at a final concentration of 1%. After incubation for 10 min, glycine was added to give a final concentration of 0.125 M for 5 min. Cells were washed twice with PBS and resuspended in Nuclei Lysis buffer (50Mm tris HCL pH 8.1; 10mM EDTA; 1%SDS and protease inhibitors) for 1 hr at +4°C. Chromatin was sonicated to obtain fragments of around 200–300 bp and then diluted 1:10 in IP Dilution buffer (0.01%SDS; 1.1% TritonX 100; 1.2mM EDTA; 16.7mM TrisHCl pH 8.1; 167mM NaCl). Chromatin extracts were immunoprecipitated overnight on rotating platform at 4°C with H3K27ac and H3K9/14ac antibodies. For each immunoprecipitation, 10 ul of antibody were used for 100ug of chromatin. Antibody-bound chromatin was incubated with 50 ul of magnetic beads (G-protein magnetic Beads, Invitrogen) 2 hrs on rotating platform at 4°C. Chromatin was washed twice with Low Salt buffer (0.1% SDS, 1% Triton, 2mM EDTA; 20mM Tris pH8, 150mM NaCl), High Salt buffer (0.1% SDS, 1% Triton, 2mM EDTA; 20mM Tris pH8, 500mM NaCl), Lithium Buffer (0.25M LiCl; 1%NP40; 1% deoxycholate; 1mM EDTA; 10mM Trish pH8) and TE. Bound DNA fragments were eluted in IP Elution Buffer (1%SDS; 1mM EDTA; 10mM Trish pH8) at 65°C for 15 minutes and the crosslink was reversed by incubation at 65°C overnight. Proteins were enzymatically digested with proteinase K, 2 hours at 37°C, and finally DNA was extracted with phenol chloroform. ChIP-seq samples were sent to IGA (Istituto di Genomica Applicata, Udine) for ChIP-sequencing on Illumina Hiseq2500 platform.

### ChIP-seq validation

ChIP assay was performed on freshly isolated FAPs following the same protocol for ChIP-seq sample preparation. Real time quantitative PCR (RT-qPCR) was performed using SYBR Green Master mix (Applied Biosystems) following manufacturer indications. Acetylation levels were normalized as percentage of input. Normal rabbit IgG (Santa Cruz Biotechnology) was used as a negative control.

Primers sequences are as follow:

CCNA2:
Fwd: GCCCTATTACCCGTCGAGTC
Rev: GTCAACCCCGAAAAACTGGC

CDK1:
Fwd: GTGAGCCTTGCCCTTCCATAA
Rev: TACCTGAGCCTGGGGACACTA

CDK4:
Fwd: GTCTATGGTCTGGCCCGAAG
Rev: CCGATCCTGGATGAGACTGC

CHEK2:
Fwd: CTCTCCGTCTCAGGAAAACTC
Rev: TTCAGCCTCATGAACTGGTAC

COX6C:
Fwd: ACGTGCAAGACATCCTAGTTC
Rev: GGTGTAGAGGTGAGAATGGTG

CYCS:
Fwd: TGATCCTAAGTGCTTCCGCC
Rev: TCTGGGAGGGTGGGTTTGTA

E2F1:
Fwd: CCCGAAGATGTCTCTAACAGTC
Rev: GGCACCAAATTCCCAATTCTG

HIST1H2AE:
Fwd: TAAAAGGCCAGACGAGAAGCT
Rev: GGAAATGAGATGTGGGAGAAGC

NDUFA4:
Fwd: AGACCGTGAACTTACGCTGG
Rev: GGAGGTCCTGGGTGACTTTG

SMAD3:
Fwd: AAAGGTTCCACATCTCAGACG
Rev: GGAAAGCAGACAAAGGAAAGTG

TGFB1:
Fwd: GGGTCGTGAGATGGAGAGAAAA
Rev: CCAACAACGTCGCTTTCCTTT

TGFB2:
Fwd: CAGCTCGGTCCTTCAGATCC
Rev: GCGAGAAAGTGCAACCTTGG

TGFBI:
Fwd: CAGAGTCAGACAAAAGTGCATG
Rev: CTGGTTTTCCCCTCTATGGTAG

TNFRSF1B:
Fwd: GCTCAGTGCCCAAAGACCTAT
Rev: AGGGGAGTAGAGTGGAAGGTG

### ChIP-seq analysis

ChIP-seq reads were aligned to the genome using bowtie-0.12.7 alignment software. Duplicated reads were removed using samtools1.3. Regions of H3K9/14ac and H3K27ac occupancy were determined using macs2 with a FDR < 0.001 and an ExtSize of 147. Input DNA of each sample was used as the control of the Peak Calling. BlackList regions of the murine genome (ENCFF547MET.bed) were excluded using bedtools software. Significant peaks revealed by macs 2 were further filtered using a threshold of macs_score > 100. Peaks were then processed with Homer to have more than 80 tags. ChIP-seq signal was visualized using NGSplot (heatmaps and average plots) and ChIP-Seeker (genomic distribution). For differential peak calling, a BedSum file, containing the regions to compare, was generated merging and sorting the two bed files of the samples of interest using bedtools. The same bed files were processed in tagDir using the Homer makeTagDirectory command. Differentially acetylated regions were generated comparing the BedSum file to either tagDir using Homer getDifferentialPeaks command with a fold change greater than 1.1.

Homer differential peaks were analysed for overlapping regions using the Intervene tool with default parameters. The matrix of overlaps was uploaded in the intervene shiny app (https://asntech.shinyapps.io/intervene/) and the Upset graph was generated.

For integration of differential ChIP-seq peaks with RNAseq, bed files generated by Homer or Intervene were intersected with a bed file containing all the promoter locations of mm10 using BedTools intersect. Peaks on the promoters were filtered in R 3.5.2 with transcripts significantly modulated (Padj < 0.1) by RNA-seq.

Bed files of differential ChIP-seq peaks and RNAseq were analysed for overlapping regions using the Intervene tool with default parameters. The matrix of overlaps was uploaded in the intervene shiny app (https://asntech.shinyapps.io/intervene/) and the Upset graph was generated. For the visualization of the genes of interest, ChIP and RNA-seq bam files were uploaded in IGV and the graphic was normalized with the group auto scale option. Coverage signal was used for ChIP-seq and FPKM value was used for RNA-seq.

### Statistical analysis

The number of independent experimental replications and precision measures are reported in the figure legends (n, mean ± sem). Comparisons between two groups were made using the student’s t-test assuming a two-tailed distribution, with significance being defined as */†p<0.05; **^/††^p<0.01; ***^/†††^p<0.001; ****^/††††^p<0.0001. Comparisons between three or more groups were made using one-way ANOVA with significance being defined as ^#^p<0.05; ##p<0.01; ^###^p<0.001; ^###^p<0.0001. Statistical analysis was conducted in PRISM 8.0 (GraphPad Software).

Significant ChIP-seq peaks were filtered for FDR < 0.001. Significant DE genes in the RNA-seq were filtered for padj<0.1.

## Acknowledgments

We thank Giovanna Borsellino and Luca Battistini (Fondazione Santa Lucia, Roma, Italy) for isolation of FAPs by cell sorting and flow cytometry analysis. This work has been supported by the following funding: AFM and DPP_Ita Grants to SC; DPP_Ita PhD fellowship to LT; R01AR076247-01 NIH/NIAMS, MDA 418870 and EPIGEN F7 to PLP.

## Author contributions

SC and LT performed most of the experimental work. SC performed ChIP-seq and RNA-seq; LT performed bioinformatic analysis; EM contributed to experimental analysis; MDB performed FACS sorting experiments; MP performed flow cytometry analysis; IS performed metabolic assays; VM, AR, SV and AM performed HDAC activity assay; SC and PLP conceived the project, supervised the study and interpreted the data. PLP wrote the manuscript and all authors discussed the results and commented on the manuscript.

---

**Supp. Figure 1.**
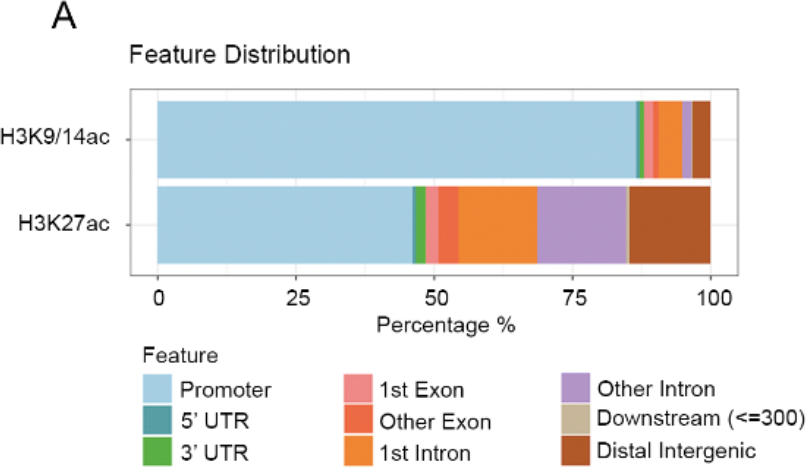
ChIP-seq genomic distribution. **A)** Graph showing the genomic distribution in term of percentage of the H3K9/14ac and H3K27ac ChIP-seq signal. On the bottom, the colorimetric legend of the features is shown.

**Supp. Figure 2.**
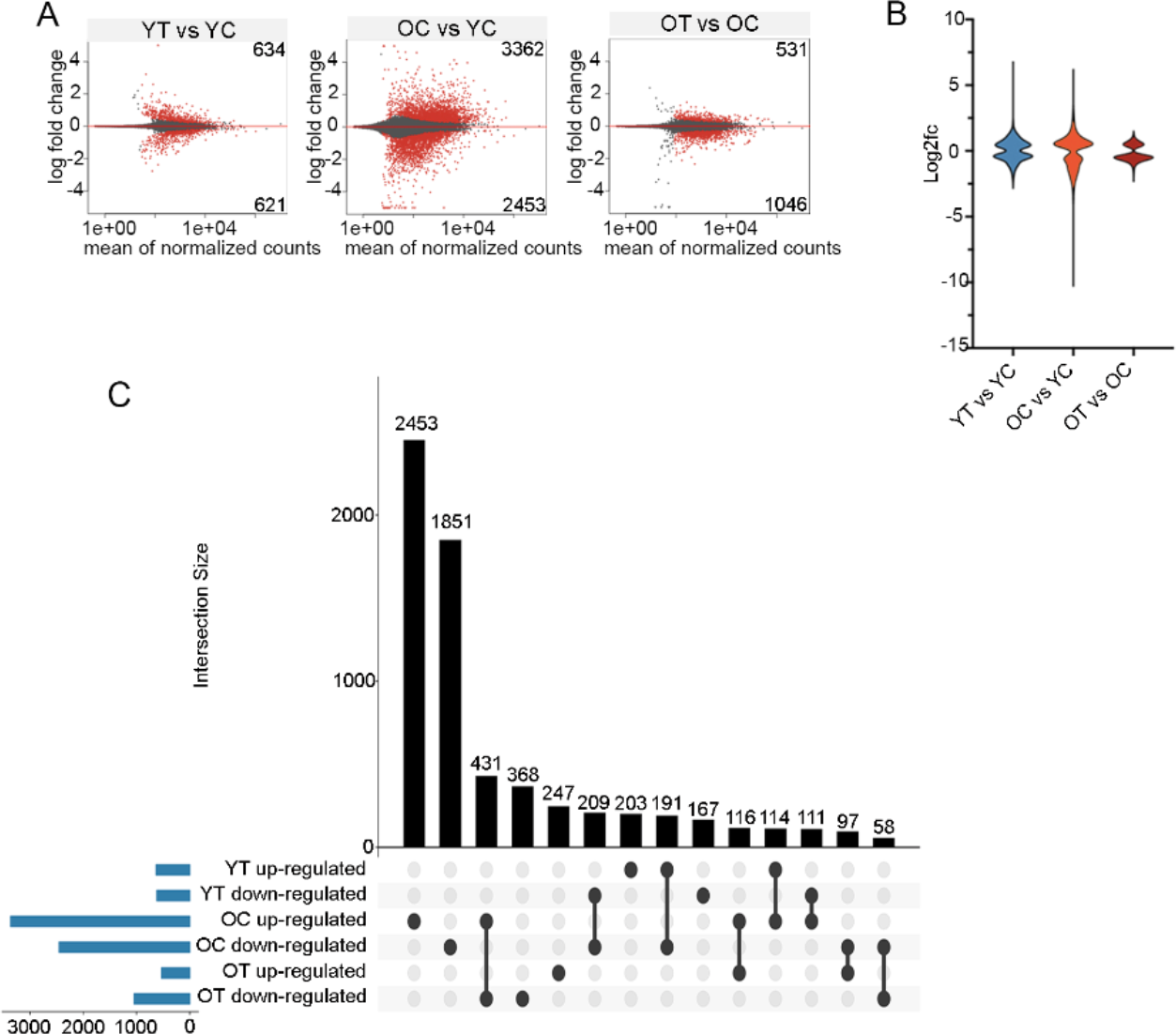
Differentially expressed genes in FAPs during disease progression and HDACi treatment. YC: young mdx FAPs; YT: young mdx FAPs *in vivo* treated with TSA for 15 days; OC: old mdx FAPs; OT: old mdx FAPs *in vivo* treated with TSA for 15 days (OT). **A)** Mean Average Plot showing the differentially expressed genes (dots, in red for padj<0.1) in the RNA-seq comparative analysis by DESEq2of young mdx FAPs treated or not with TSA (left panel), young and old mdx FAPs (middle panel) old mdx FAPs treated or not with TSA (right panel). **B)** Violin plot showing the distribution of the log_2_ fold change in the same analysis shown in A). **C)** Upset graph showing the intersection size (in black) between the differentially expressed genes (in blue) in the same experimental conditions described in A). The top 14 intersections are shown.

**Supp. Figure 3.**
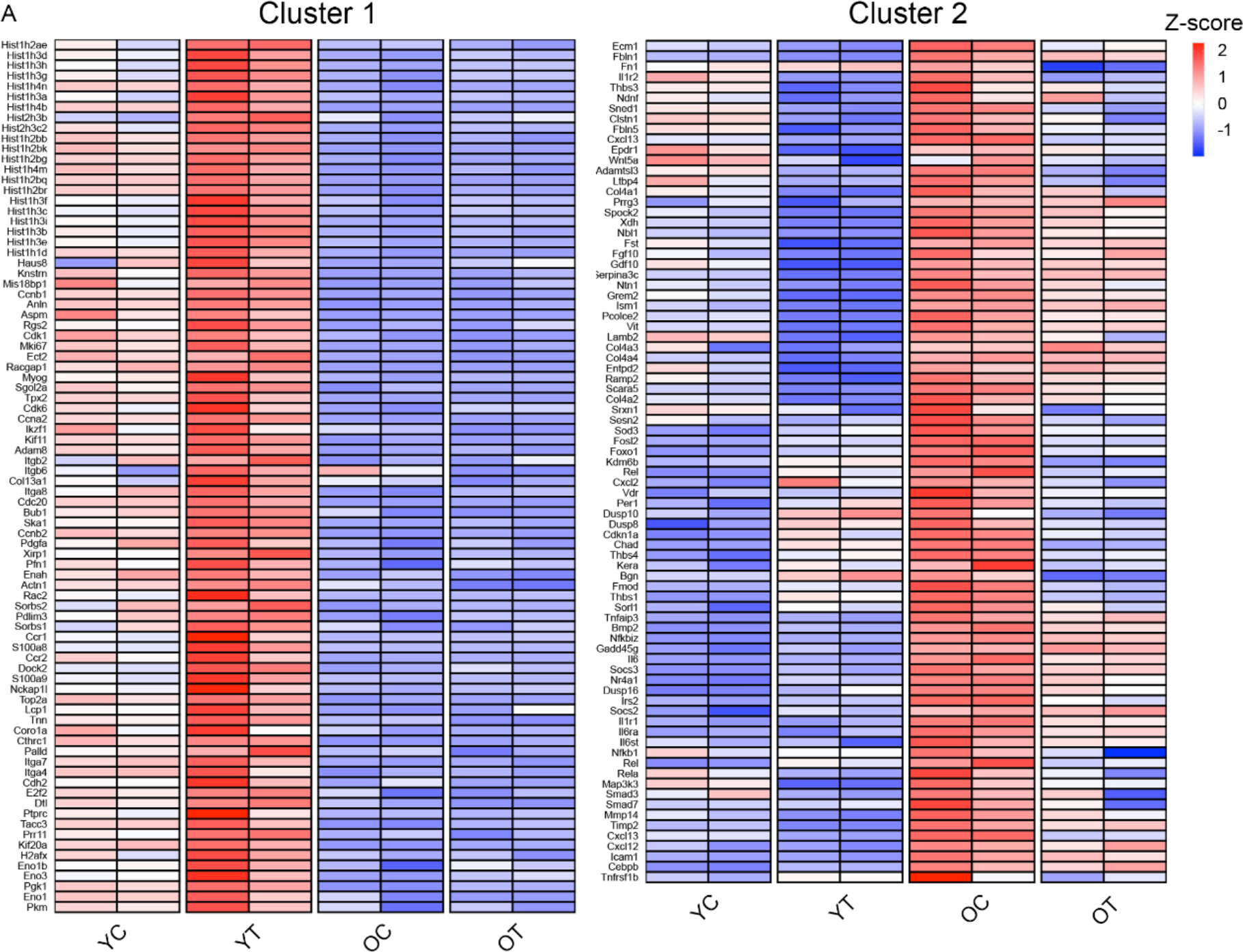
Cluster 1 and 2 gene list. YC: young mdx FAPs; YT: young mdx FAPs *in vivo* treated with TSA for 15 days; OC: old mdx FAPs; OT: old mdx FAPs *in vivo* treated with TSA for 15 days (OT). **A)** Heatmap showing the name of cluster 1 and cluster 2 DE genes. Gene expression is represented as z-score calculated across the experimental conditions.

**Supp. Figure 4.**
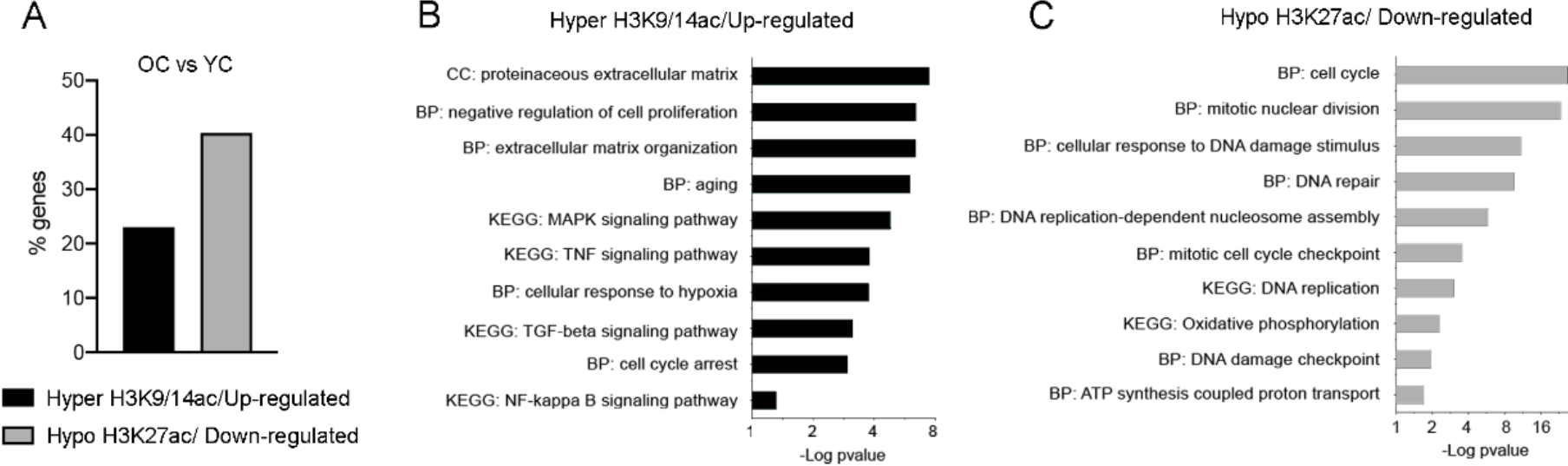
ChIP and RNA-seq integration in old mdx FAPs. YC: young mdx FAPs; OC: old mdx FAPs. **A)** Graph showing the percentage of genes found in the integration between differentially expressed (RNA-seq) and differentialy acetylated (ChIP-seq) genes in the comparison between young and old mdx mice. **B)** Gene Ontology performed on genes both hyper-acetylated in H3K9/14 and up-regulated in old mdx FAPs versus young mdx FAPs. **C)** Gene Ontology performed on genes both hypo-acetylated in H3K27 and down-regulated in in old mdx FAPs versus young mdx FAPs.

**Supp. Figure 5.**
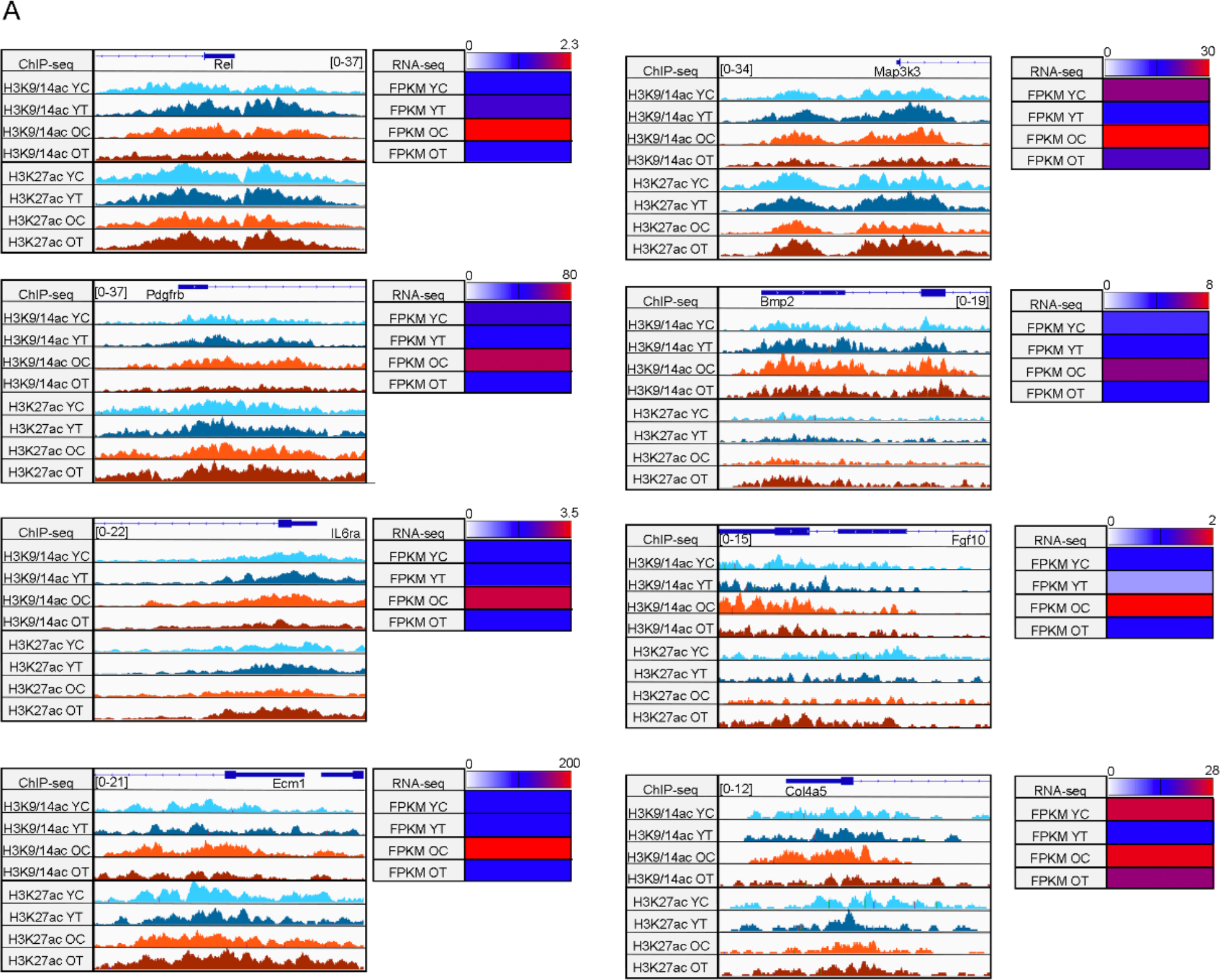
Acetylation and expression profile of SASP genes. YC: young mdx FAPs; YT: young mdx FAPs *in vivo* treated with TSA for 15 days; OC: old mdx FAPs; OT: old mdx FAPs *in vivo* treated with TSA for 15 days (OT). **A)** Acetylation tracks for H3K9/14ac and H3K27ac ChIP-seq (on the left) and corresponding FPKM values by RNA-seq (on the right) for representative genes involved in SASP.

**Supp. Figure 6.**
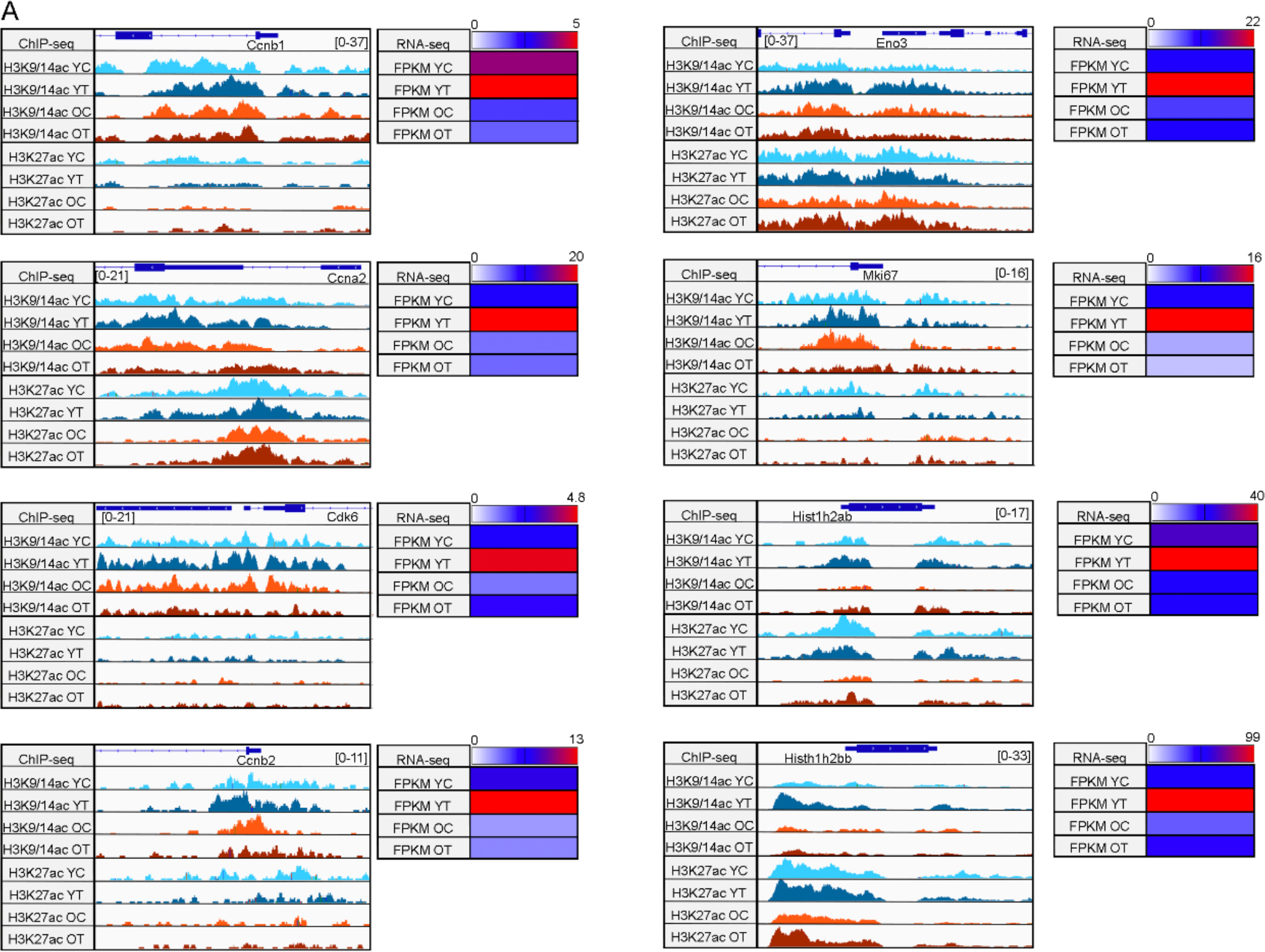
Acetylation and expression profile of cell cycle genes. YC: young mdx FAPs; YT: young mdx FAPs *in vivo* treated with TSA for 15 days; OC: old mdx FAPs; OT: old mdx FAPs *in vivo* treated with TSA for 15 days (OT). **A)** Acetylation tracks for H3K9/14ac and H3K27ac ChIP-seq (on the left) and corresponding FPKM values by RNA-seq (on the right) for representative genes involved in cell cycle activation.

**Supp. Figure 7.**
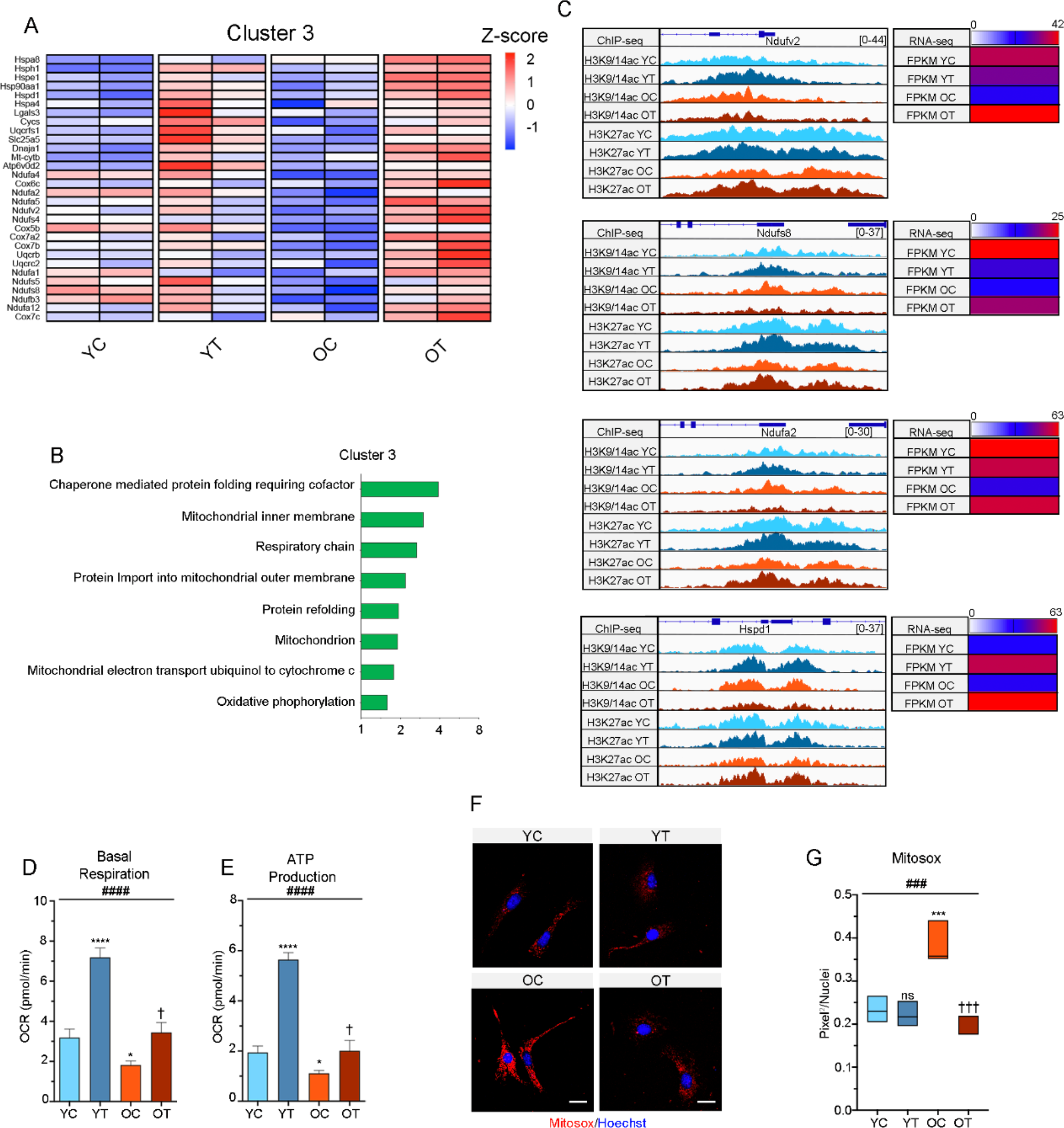
Acetylation and expression profile of genes involved in mitochondrial function. YC: young mdx FAPs; YT: young mdx FAPs *in vivo* treated with TSA for 15 days; OC: old mdx FAPs; OT: old mdx FAPs *in vivo* treated with TSA for 15 days (OT). **A)** Heatmap showing the name of cluster 3 DE genes. Gene expression is represented as z-score calculated across the experimental conditions. **B)** Gene Ontology performed on cluster cluster 3 DE genes. **C)** Acetylation tracks for H3K9/14ac and H3K27ac ChIP-seq (on the left) and corresponding FPKM values by RNA-seq (on the right) for representative genes involved in mithocondrial function. **D)** Graphs showing the Basal Respiration measured as oxygen consumption rate (OCR) by Seahorse in young and old mdx FAPs treated or not with TSA. **E)** Graphs showing the levels of ATP production measured as OCR after oligomycin administration by Seahorse in the same experimental points described in D). **F)** Representative images of the Mitosox staining (red mitochondrial superoxide indicator) in young and old mdx FAPs treated or not with TSA. Nuclei were counterstained with Hoechst 33342 (Hoechst, in blue); Scale bar = 25 μm; **G)** Box plot showing the quantification of the red Mitosox staining in the same experimental conditions described in H). Data are represented as average ± SEM (n=3). *p<0.05, ***p<0.001, ****p<0.0001 against young mdx FAPs by t-test; ^†^p<0.001, ^†††^p<0.001 against old mdx FAPs by t-test; ^###^p<0.001, ####p<0.0001 via one way ANOVA.

